# Structure of the connexin-43 gap junction channel in a putative closed state

**DOI:** 10.1101/2022.03.26.485947

**Authors:** Chao Qi, Silvia Acosta-Gutierrez, Pia Lavriha, Alaa Othman, Diego Lopez-Pigozzi, Erva Bayraktar, Dina Schuster, Paola Picotti, Nicola Zamboni, Mario Bortolozzi, Francesco L. Gervasio, Volodymyr M. Korkhov

**Affiliations:** Institute of Molecular Biology and Biophysics, ETH Zurich, Switzerland; Laboratory of Biomolecular Research, Paul Scherrer Institute, Villigen, Switzerland; Institute for the Physics of Living Systems, Institute of Structural and Molecular Biology, and Department of Chemistry. University College London, London, UK; Institute for Bioengineering of Catalunya (IBEC), The Barcelona Institute of Science and Technology, Barcelona, Spain; Institute of Molecular Systems Biology, ETH Zurich, Switzerland; Department of Physics and Astronomy “G. Galilei”, University of Padova, Padua, Italy; Veneto Institute of Molecular Medicine (VIMM), Padua, Italy; Department of Chemistry, University College London, WC1E 6BT London, UK; School of Pharmaceutical Sciences, University of Geneva, CH-1211 Geneva, Switzerland; ISPSO, University of Geneva, CH-1211 Geneva, Switzerland

## Abstract

Gap junction channels (GJCs) mediate intercellular communication by connecting two neighboring cells and enabling direct exchange of ions and small molecules. Cell coupling via connexin-43 (Cx43) GJCs is important in a wide range of cellular processes in health and disease ^1-3^, yet the structural basis of Cx43 function and regulation has not been determined until now. Here we describe the structure of a human Cx43 GJC solved by cryo-EM and single particle analysis at 2.26 Å resolution. The pore region of Cx43 GJC features several lipid-like densities per Cx43 monomer, located close to a putative lateral access site at the monomer boundary. We found a previously undescribed conformation on the cytosolic side of the pore, formed by the N-terminal domain and the transmembrane helix 2 of Cx43 and stabilized by a small molecule. Structures of the Cx43 GJC and hemichannels in nanodiscs reveal a similar gate arrangement. The features of the Cx43 GJC and hemichannel cryo-EM maps and the channel properties revealed by molecular dynamics simulations suggest that the captured states of Cx43 are consistent with a closed state.

## Introduction

Gap junction (GJ) mediated intercellular communication is one of the major pathways of information exchange between the cells. GJs are specialized regions of the plasma membrane at the cell-cell interface that link two adjacent cells and establish their metabolic and electrical coupling ^4^. Connexins are the building blocks of the GJ channels (GJCs) which belong to a group of large-pore channels. This group includes a number of structurally related (innexins, pannexins, LRRC8) and unrelated proteins (CALHM) ^5^. A total of 21 connexin genes have been identified in the human genome ^6^. The 43 kDa connexin-43 (Cx43, gene name *GJA1*) was identified as a major constituent of rat heart GJs in 1987 ^7^, and it is arguably one of the most extensively studied connexins. Like all connexin homologues, Cx43 monomers assemble into hexameric hemichannels (HCs), also known as connexons. HCs that reach the plasma membrane of one cell may interact with their counterparts on the neighboring cell, forming GJCs, typically organized into hexagonal arrays at the intercellular interface ^8^. GJCs enable direct metabolic and electric coupling between the cells, facilitating the passage of ions, small molecules, metabolites, peptides and other cellular components below a size threshold of approximately 1.5 kDa ^9^. GJCs formed by Cx43 are crucial for a wide range of physiological processes, from propagation of heart action potentials ^3^ to maintenance of neuro-glial syncytium ^2^ and skin wound healing ^1^. The clinical importance of Cx43 is highlighted by mutations linked to several genetic disorders, such as oculodentodigital dysplasia (ODDD) ^10-12^, hypoplastic left heart syndrome 1 ^13^, Hallermann-Streiff syndrome ^14^, atrioventricular septal defect 3 ^13^, etc., and by recognition of the protein as a drug target for treatment of cancer, skin wounds, eye injury and inflammation and cardiac arrhythmias (reviewed by Laird and Lampe ^9^).

Much of what we know today about the molecular biology, electrophysiological properties and regulation of connexin gap junctions has been derived from the studies on Cx43 (reviewed in ^15^). Early attempts to characterize the structure of Cx43 using cryo-electron microscopy (cryo-EM) of 2D crystals produced low resolution reconstructions ^16^. More recently, several homologous connexin GJCs and HCs have been structurally characterized at high resolution, using X-ray crystallography and cryo-EM. The structures of Cx26 ^17^ and Cx46/50 GJCs ^18,19^, together with the recent structure of Cx31.3 HC ^20^ provided deep insights into the shared structural features of the connexin channels. These structures hint at the role of the N-terminal domain (NTD) in molecular gating of GJCs and HCs, with several conformations of the NTD observed in different connexin homologues ^18-20^. Additionally, biochemical evidence points to the roles in channel gating played by the link between Cx43 intracellular loop and the C-terminal region ^21,22^. However, despite the availability of this evidence, the molecular determinants of intracellular connexin channel gating remain unclear. To shed light on the structural basis of Cx43 gating, we set out to determine its structure by cryo-EM and to analyse its dynamics with molecular dynamics (MD) simulations.

## Results

### Structures of Cx43 GJC and HC in detergent micelles and in nanodiscs

Our Cx43 expression system was tested using electrophysiology in HeLa cells (Extended Data Fig. 1a-d). The experiments confirmed the functionality of Cx43 GJCs expressed in transfected cells. For large-scale protein expression, Cx43 featuring a C-terminal HRV3C-YFP-twinStrep tag, was expressed in adherent mammalian cells (HEK293F) using transient transfection method. The produced protein was purified using affinity and size exclusion chromatography in digitonin (Extended Data Fig. 2a-c) and flash frozen on cryo-EM grids. The coomassie-stained SDS PAGE gel bands of the purified protein (Extended Data Fig. 2c) are consistent with the western blot analysis (Extended Data Fig. 2d) of the expressed untagged Cx43. The grids were subjected to single particle cryo-EM analysis (Extended Data Fig. 3), yielding the final 3D reconstruction at 2.26 Å resolution (Fig. 1a, Extended Data Fig. 3-4).

**Figure 1.**
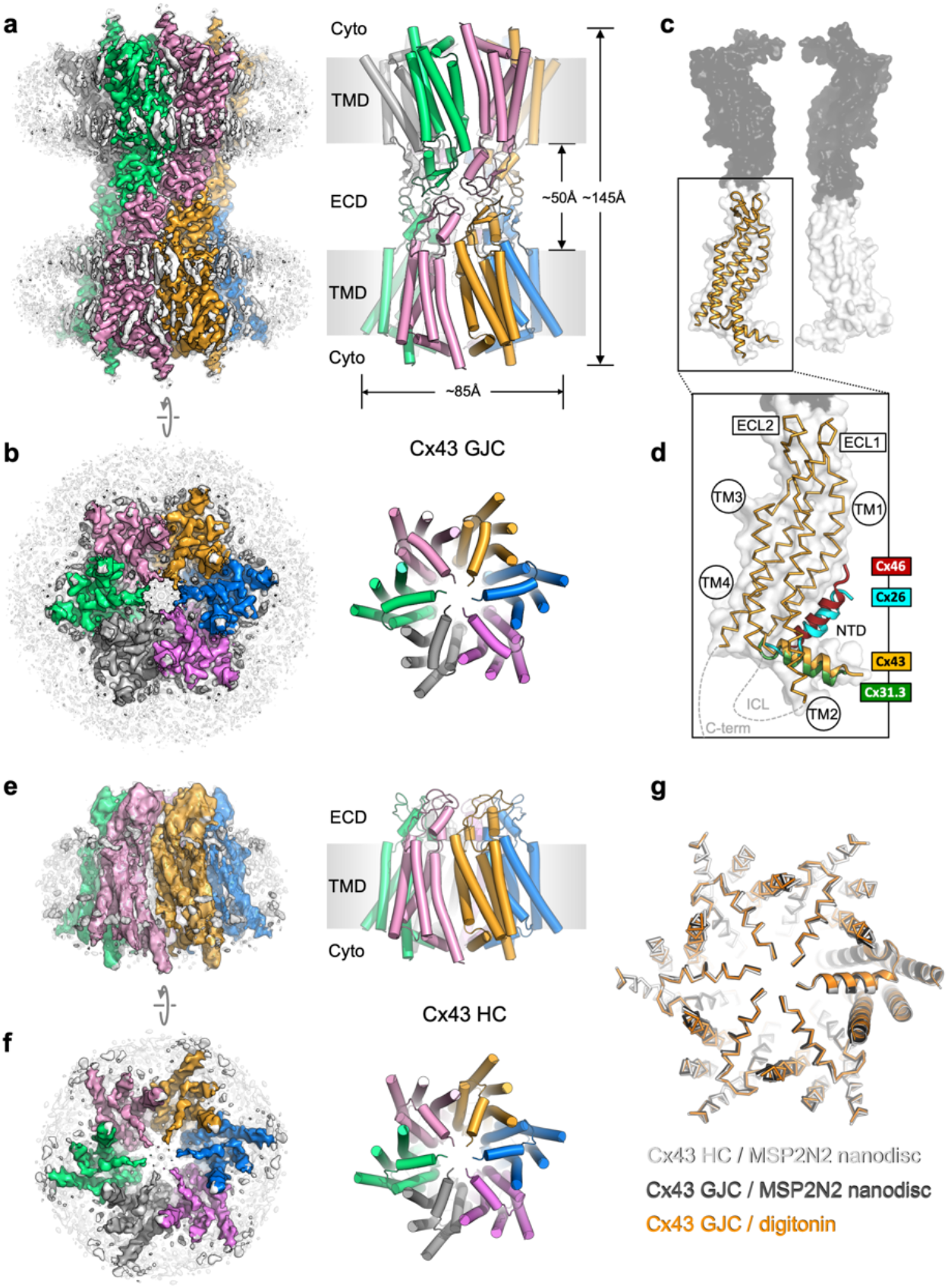
Structure of human Cx43 gap junction channel. **a-b**, Cryo-EM density map and model of Cx43 GJC solved by cryo-EM at 2.26 Å resolution. The individual Cx43 monomers in each HC within the GJC are coloured blue, pink, grey, green, salmon and orange. Grey densities correspond to the detergent micelle and the bound sterol-like molecules. **c**, The position of a single Cx43 monomer (orange) within a GJC (represented as a surface of juxtaposed Cx43 monomers in two distinct membrane regions, white and grey). **d**, Alignment of the monomers of Cx43, Cx26 (PDB ID: 2zw3), Cx46 (PDB ID: 7jkc) and Cx31.3 (PDB ID: 6l3t) shows the distinct orientations of the N-terminal domain (“NTD”) helix. Individual TM domains, extracellular loops 1 and 2 (“ECL1-2”), relative positions of the intracellular loop (“ICL”) and the C-terminus (“C-term”) are indicated. **e-f**, Same as a-b, for the Cx43 HC in MSP2N2 nanodiscs at 3.98 Å resolution. **g**, Alignment of three indicated structures shows that the conformation of the gating region is highly conserved. The models are shown as ribbons; one of the aligned monomers in each of the structures is represented as cartoon, to clearly indicate the monomer boundaries.

The overall architecture of the protein complex resembles that observed with other connexin GJCs, with two hemichannels (connexons) from adjacent plasma membrane regions coupled to form a full GJC (Fig. 1a-b). The exterior of the channel is decorated by several ordered detergent and/or lipid-like density elements (Fig. 1a-b), reminiscent of the previously observed lipids bound at the protein-bilayer interface in other connexins ^19^. In the case of Cx43, the lipid-like densities appear to decorate the protein-lipid bilayer interface at both the inner and the outer leaflet of the membrane. Comparison of the Cx43 monomers to the available structures of Cx26 and Cx46 GJCs and the Cx31.3 HC revealed a major difference in the NTD arrangement in these channels (Fig. 1c-d). The conformation of the NTD in the Cx31.3 HC structure appears to be the closest to that in Cx43 GJC. Analysis of the tryptic peptides revealed that most of the protein lacks the residue M1 (Extended Data Fig. 5), and thus the model was built starting with G2.

To ascertain that the conformation of Cx43 GJC is not induced by the detergent present in the sample, e.g. through detergent binding at specific sites on the protein surface, we removed the detergent and reconstituted the protein into MSP2N2 nanodiscs, using 1-palmitoyl-2-oleoyl-glycero-3-phosphocholine (POPC) as a mimic for the native lipid environment. The reconstituted protein was subjected to the same imaging and analysis workflow as Cx43 GJC in digitonin. The 2D classes of the MSP2N2-reconstituted Cx43 showed features consistent with a mixture of GJCs and HC. Processing of the corresponding particles resulted in 3D reconstructions of the Cx43 GJC at 2.95 Å resolution (Extended Data Fig. 6, 8) and HC in nanodiscs at 3.98 Å resolution (Fig. 1e-f, Extended Data Fig. 7, 8). The GJCs in detergent and in nanodiscs are nearly identical, with an RMSD of 0.97 Å between the aligned Cx43 monomers (Fig. 1g, Extended Data Fig. 9a). Although the differences between the HC in nanodisc and the GJC are more pronounced (Extended Data Fig. 9a), the conformation of the cytosolic region and the NTD is highly conserved (Fig. 1g).

### Conformation of the Cx43 putative gate region

The reconstructed putative gate region of Cx43 shows several narrow openings connecting the pore vestibule and the pore interior: a single ∼6-7 Å wide central opening and six adjacent openings of similar dimensions (Fig. 2a-b). This arrangement of the gate region is distinct from any of the previously observed states of connexin GJCs or HCs. This particular gate conformation is established through an interplay of two structural elements: the NTD and the TM2 (Fig. 2c-d). The NTD of Cx43 is arranged near parallel to the membrane plane, in a centre-oriented conformation (Fig. 2c). The HC of Cx31.3 assembles in a similar manner, with a well-ordered NTD resolved by cryo-EM (Fig. 2c-d). However, the TM2 region of Cx31.3 forms a tight seal with the NTD (Fig. 2e). In contrast, the TM2 of Cx43 is shifted away from the pore centre, creating six openings (Fig. 2f). Alignment of the protomers of Cx43 and Cx31.3 showed an RMSD of 1.78 Å (using cealign in PyMol). In contrast, the RMSD value for the region corresponding to the NTD and TM2 was 5.675 Å (calculated using rms_cur in PyMol, with the residue range selections of 2-47 and 2-45 for the aligned protomers of Cx43 and Cx31.3, respectively), further confirming that these two regions differ substantially between Cx43 and Cx31.3. The arrangement of the Cx43 NTD appears to be stabilized by a small molecule: a density element likely corresponding to a bound small molecule is present within this region (referred to as the “NTD lipid site”), wedged between the adjacent NTDs (Fig. 2g). Thus, although the conformation of the Cx43 gate features an opening, this site is blocked by a yet unidentified small molecule (which we refer to as “lipid-N”).

**Figure 2.**
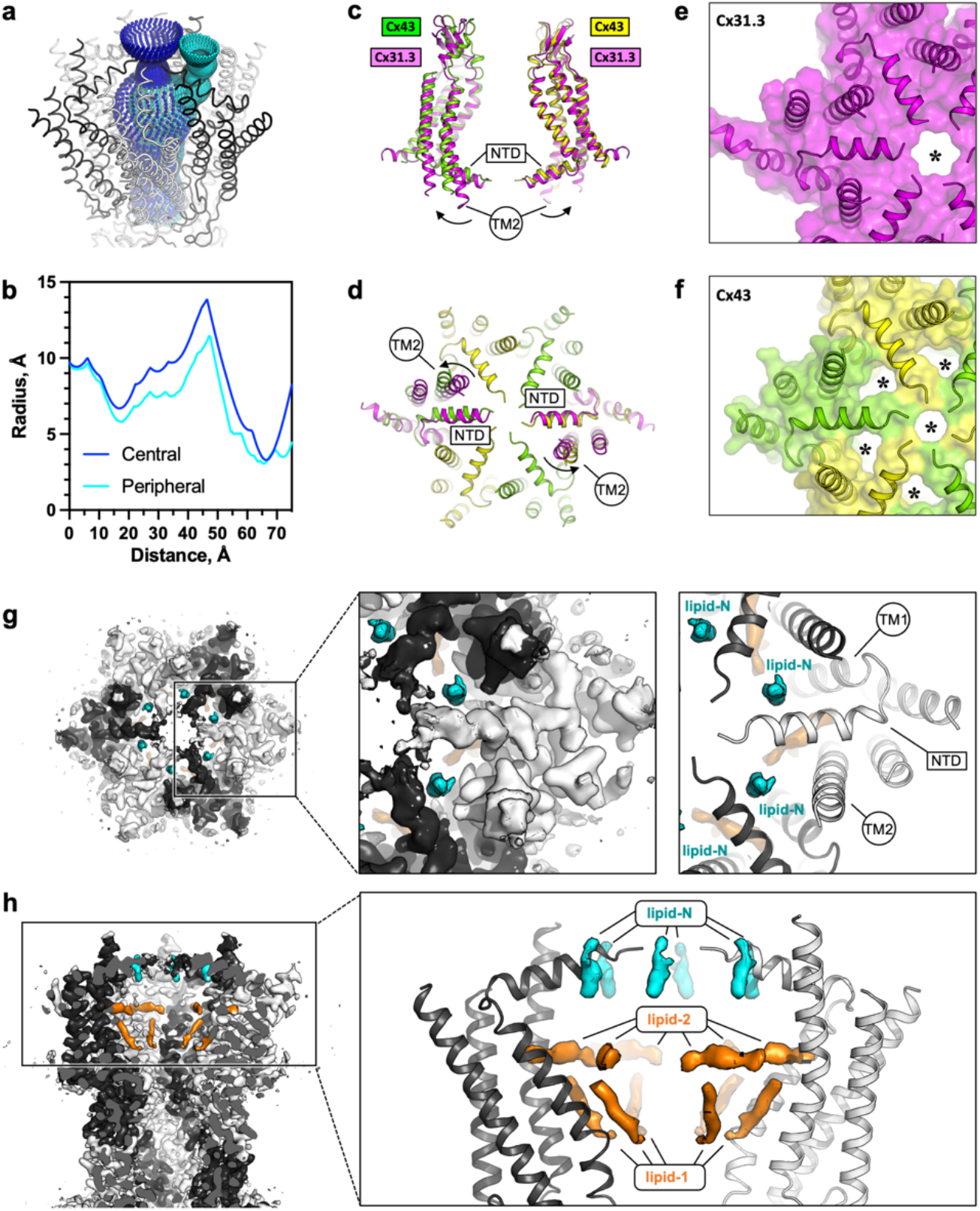
The Cx43 gate adopts a closed conformation. **a-b**, Analysis of the pore opening dimensions using HOLE reveals a constriction of the pore in the putative gate region. Only a central opening and one of the six peripheral openings within one HC of Cx43 GJC in digitonin is shown. The distance is calculated from the center of the GJC pore to a point outside of the channel. Central and peripheral openings are coloured blue and cyan, respectively. **c-d**, Comparison of the Cx43 in digitonin with the Cx31.3 HC shows that the peripheral opening is created by the particular arrangement of the NTD and by adjustment of the TM2. **e-f**, The distinct NTD/TM2 arrangement results in a single pore opening in the structure of Cx31.3 (indicated with an asterisk, e), contrary to Cx43 (f). **g**, A view of the Cx43 gating region from the cytosol reveals the location of the “lipid-N” molecules stabilizing the NTD arrangement, shown as isolated densities (cyan). **h**, A slab view of the gating region parallel to the membrane plane shows the relative arrangement of the lipid-N and the intra-pore lipid densities (“lipid-1” and lipid-2”; orange).

While a direct comparison of the Cx43 gating regions of the Cx43 and Cx31.3 HCs is possible (Extended Data Fig. 9c), the recently determined 3D reconstruction of the Cx26 mutant N176Y was not accompanied by a new atomic model ^23^. Instead the study reporting this reconstruction compared the density map with an HC model from a Cx26 GJC structure (PDB ID: 5ERA) ^24^. As this model lacks the NTD, any comparisons of the gating region in the Cx43 HC with that in the Cx26 HC are presently limited (Extended Data Fig. 9d).

In addition to the lipid-N molecules stabilizing the NTD arrangement, the cryo-EM density of Cx43 GJC features several well resolved densities inside the pore region (lipid-1 and lipid-2, Fig. 2h). These elements likely correspond to bound sterol molecules, such as cholesterol co-purified with the protein from the mammalian cells or cholesterol hemisuccinate (CHS) added to the solubilization mixture during protein extraction from the membrane. Similar densities are present in the Cx43 GJC reconstruction in nanodiscs, in the absence of detergent molecules (Extended Data Fig. 10a). Additionally, it is noteworthy that the annular lipid densities are conserved in both the detergent-solubilized and the nanodisc-reconstitued Cx43 GJCs, indicating that the protein-lipid interface of Cx43 features sites where ordered lipid molecules may bind (Extended Data Fig. 10b). The functional significance of the ordered annular lipids for the channel activity and for GJ plaque assembly remains to be carefully investigated.

Identification of the lipid-like molecules bound to Cx43 is a significant challenge. Despite the high overall resolution of the 3D reconstruction of the Cx43 GJC (2.26 Å resolution in detergent and 2.95 Å in nanodisc; Extended Data Fig. 3, 6), the cryo-EM map features in regions corresponding to the lipid densities are insufficient to assign the lipid identity unambiguously. To determine the identities of these lipids experimentally, we performed lipidomic analysis of the organic extracts prepared from the purified Cx43 samples and compared them to the mock control (extracts of prepared from eluates of the purification procedure using cells that do not overexpress the protein as starting material). The results showed that several types of phospholipids, as well as cholesterol were present in both samples. The only lipid-like compound specifically enriched in the purified sample was dehydroepiandrosterone (DHEA; Extended Data Fig. 10c). DHEA is a most abundantly expressed neurosteroid known to modulate ligand-gated ion channels, such as GABA_A_ ^25^ or NMDA and AMPA receptors ^26^. Although we are presently not able to conclusively state that DHEA corresponds to any of the densities present in our reconstruction, its enrichment in the sample is an interesting observation that deserves detailed future investigations. While the density corresponding to lipid-N may correspond to DHEA (Extended Data Fig. 4e), it is also possible that this density corresponds to a larger molecule that is only partially ordered.

### Cx43 junction interface

As in other connexins, the Cx43 GJC interface is formed by two extracellular loops, ECL1 and ECL2 (Fig. 3a-b). The portion of the ECL1 comprising residues N55-Q58 makes contacts with two Cx43 monomers from the opposite membrane, with N55-Q58 of one monomer and Q58’-P59’ of the neighboring monomer within 4 Å distance (Fig. 3c, *top left*). The ECL2 region that directly participates in sealing the junction includes residues P193-D197. Comparisons of the junction-forming loops in Cx43 to those in Cx46 (Fig. 2c, *middle*) and Cx26 (Fig. 2c, *right*) show that each protein uses a similar pattern of inter-HC interaction within each GJC. It is noteworthy that both the amino acid sequences (Fig. 3d) and the interaction networks (Fig. 3c) are similar in the ECL1 for Cx43 and Cx46, and in the ECL2 for Cx46 and Cx26. The differences in these two loops, and especially in ECL2, underlie the inability of Cx43 (which has been grouped into a docking group 2) to engage in heterotypic interactions with group 1 connexins (such as Cx46 and Cx26) ^27^. Our structure is thus in line with the recognized consensus motif necessary for heterotypic complementarity of connexin GJCs ^28^.

**Figure 3.**
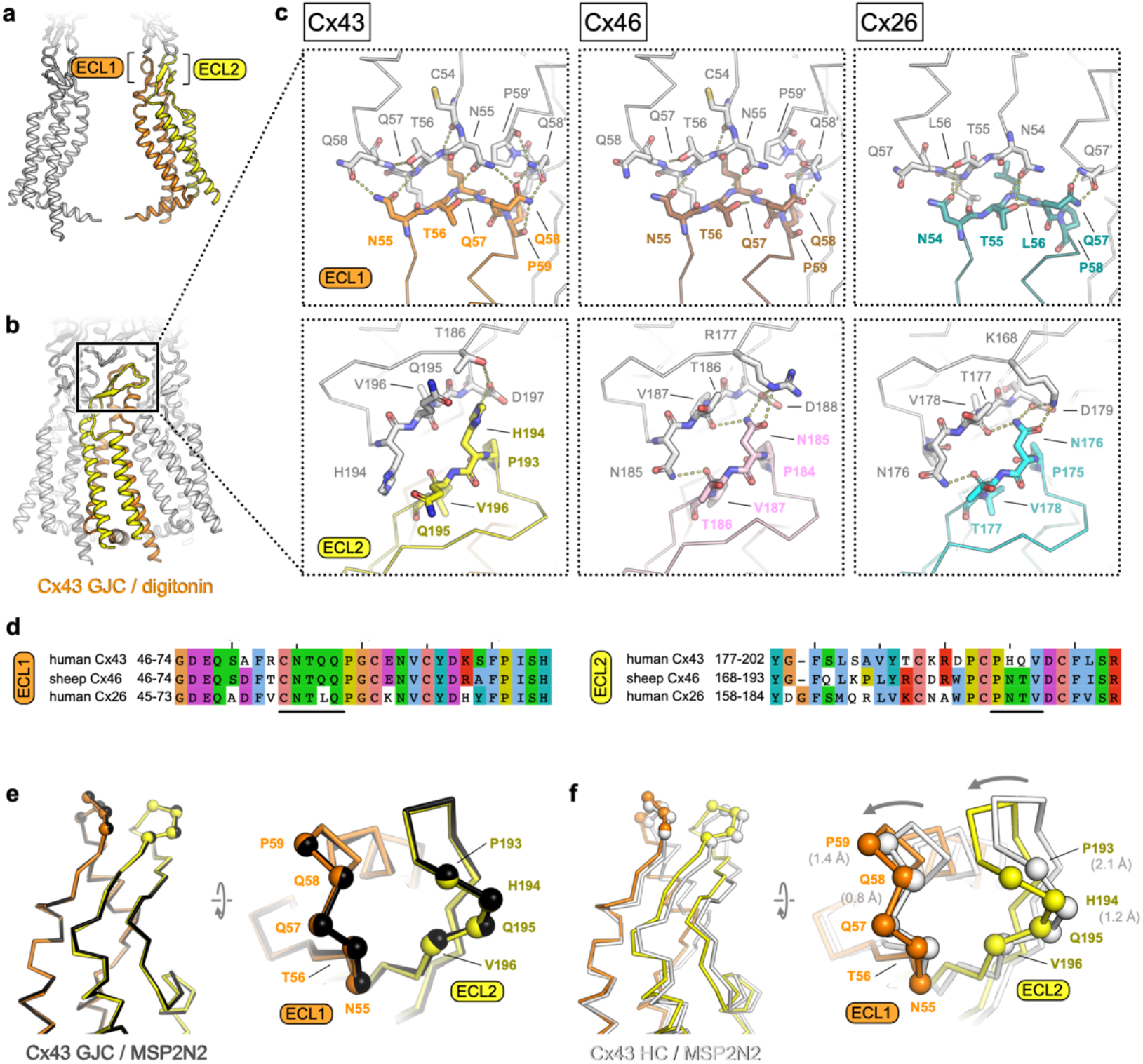
Extracellular domain of Cx43 GJC and HC. **a-b**, The lateral views of Cx43 GJC in digitonin, indicating the positions of the extracellular loops 1 and 2. **c**, Views of the extracellular loops ECL1 (top) and ECL2 (bottom), for Cx43 (left), Cx46, (middle; PDB ID: 7jkc), and Cx26 (right; PDB ID: 2zw3). The residues of one monomer in each of the structures, directly involved in junction formation, are coloured by element with carbon atoms as orange/yellow (Cx43), brown/pink (Cx46) and teal/cyan (Cx26). The residues of the neighbouring connexin monomers within 4 Å distance are shown coloured with white carbon atoms. Dotted lines indicate electrostatic contacts, calculated in PyMol. **d**, Sequence alignment of the complete ECL1 and ECL2 regions of the three proteins shown in c. The black lines indicate the GJC interface residues, as shown in c. **e**, Alignment of the Cx43 GJC structures in digitonin micelles and in nanodiscs. Cα atoms of the interface residues shown in C are represented as spheres (the ribbon and spheres of Cx43 in nanodiscs is coloured black). **f**, Same as e, for a comparison between Cx43 GJC in digitonin and Cx43 HC in nanodiscs. The arrow indicates the movement of the two loops that accompanies GJC formation. The ribbon and spheres of Cx43 HC are light grey. The grey numbers in brackets indicate displacement of selected Cα atoms (residues P59, Q58, P193, P194).

### Conformation of the Cx43 hemichannel

Although the two Cx43 GJC structures (in detergent micelles and in nanodiscs) are almost identical, the Cx43 HC shows a notable difference in the ECL1-2 conformation (Fig. 3e-f). The resolution of the HC reconstruction does not allow us to make definitive statements about the orientations of individual side chains, but it is clear that both loops engaging in junction formation move inward upon docking of the two HCs (Fig. 3f). This conformational change likely involves intra- and intermolecular cooperativity. The ECL1 and ECL2 are linked via several disulphide bonds, and thus the rearrangement within each molecule would require concerted movement of the whole extracellular domain (ECD). The ECD movement within one monomer is likely cooperatively coupled to the neighboring chains within the Cx43 HC.

### Disease-linked mutations in Cx43

A number of disease-linked mutations associated with Cx43 can be mapped directly to three regions of interest revealed by our structures: (i) the GJC intra-pore lipid-like site, (ii) the ECD, and (iii) the gating region (Fig. 4a). The interior of the channel features two lipid-like densities (Fig. 2h, 4b; Extended Data Fig. 4e, 10a). The observed position of lipid 1 is consistent with the previously found phospholipid-like densities inside the pore of the Cx31.3 HC ^20^. Lipid 2 inserts into the pocket formed by TM1 and TM2, parallel with the NTD. Unlike Cx31.3, where elongated densities within the pore region could be interpreted as hydrophobic tails of bound phospholipids, the density in the Cx43 GJC appears consistent with that of a sterol (Fig. 2h, 4b). The presence of a small hydrophobic small molecule in Cx43 at this site suggests a potential mechanism of lipid-based Cx43 regulation, whereby binding of a lipid could directly influence the conformation of the gating elements of the protein (such as NTD). An effect of a cholesterol analogue 7-ketocholesterol on Cx43 permeability has been observed previously ^29^, and such an effect may be mediated via the binding sites within the GJC (or HC) pore (Fig. 4b; Extended Data Fig. 10a). The presence nearby of several residues known to be linked to ODDD when mutated (S27P, I31M^30^, S86Y^31^, L90V^10^) points to a potential functional significance of the intra-pore lipid binding site.

**Figure 4.**
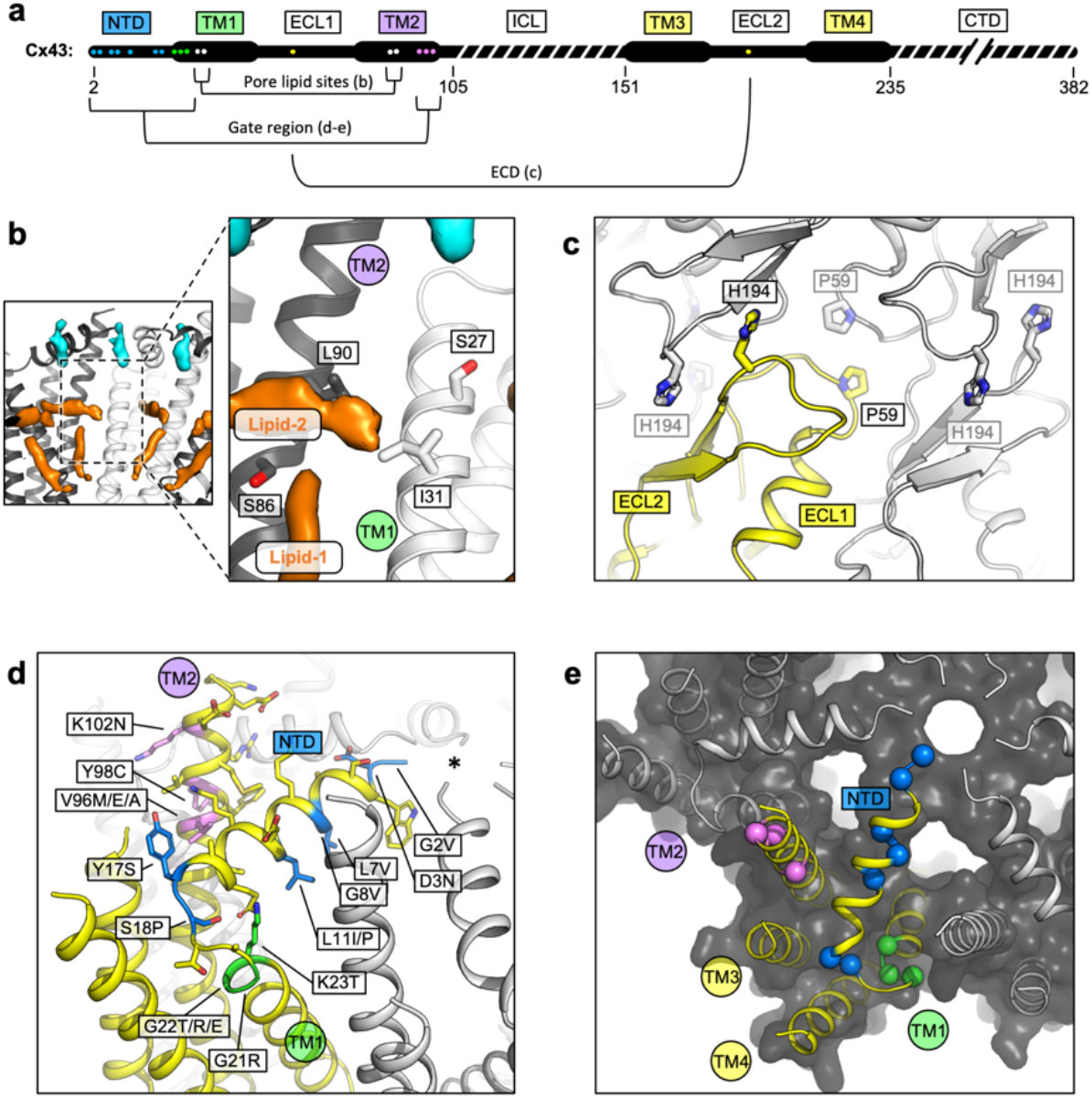
Locations of the disease-linked mutations in the Cx43 GJC structure. **a**, Disease-linked mutations mapped on the sequence of Cx43. The amino acid residues proximal to the intrapore lipid densities (“Pore lipid sites”) are shown as white dots. The residues at the GJC interface are shown as yellow dots (“ECD”). The residues of the putative gating region (“Gate region”) are shown as blue (NTD), green (TM1) and violet (TM2) dots. The sequence elements not resolved in our 3D reconstruction are indicated with a dashed line. **b**, Two sterol-like density elements (lipid-1 and -2) are located within the pore region of Cx43 GJC (white). The indicated disease-linked mutations are located within close distance of the pore lipid densities. **c**, A view of the extracellular loops forming the junction, indicating two residues known to be linked with ODDD, P59 and H194 (side chains shown as sticks; one of the monomers in the GJC is coloured yellow). **d**, A view of the gating region with the highlighted disease-linked mutations, colored as in a. **e**, A similar representation of the disease-linked mutations, with Cα atoms shown as spheres, illustrating the contribution of the mutants to gate formation.

Two known mutations linked to ODDD are located in the Cx43 ECD: P59 in ECL1 ^32^ and H194 in ECL2 ^33^. As suggested by the Cx43 GJC structures, these residues are located in conserved regions where any substitution can be expected to cause disruption of the contacts critical for junction formation (Fig. 4c).

Multiple mutations in Cx43 associated with ODDD are located in the N-terminus (G2V, D3N, L7V, L11I/P, Y17S, S18P), the TM1 region (G21R, G22E, K23T) and the TM2 region proximal to the gate (V96M/E/A, Y98C, K102N) ^11^. In some cases, the mutations reduce the ability of Cx43 to form the gap junction plaques at the plasma membrane, as is the case for Y17S or G21R (as well as L90V, located at the pore lipid binding site) ^34^. The mutations tend to have deleterious effects on the permeability of Cx43 to ions and small molecules ^11^. The mutations G2V, D3N, W4A and L7V were shown to eliminate the function of Cx43 GJCs ^35^. Interestingly, a G8V amino acid substitution has been linked to palmoplantar keratoderma and congenital alopecia 1 (PPKCA1). This mutant can form functional gap junctions and has enhanced HC activity ^36^. Mapping these sites on the structure of Cx43 allows us to gain insights into the possible mechanisms that underlie the associated disorders (Fig. 4d-e). For example, channel blockage or hyperpermeability due to mutations in or around the gating region may link Cx43 to diseases.

### Molecular dynamics simulations of Cx43 GJC

Based on our model of Cx43, the dimensions of the pore opening are likely incompatible with the translocation of a larger molecule, such as ATP, cAMP or IP3 (typical substrates of Cx43 GJC-mediated transport). To assess the permeability of the gate to ions, we performed MD simulations. We performed eighteen independent molecular dynamics simulations of the Cx43 GJC embedded in a double POPC bilayer solvated in 150 mM KCl. Given the ambiguity in the lipid identity and binding mode, in the first set of simulations, no potential lipid surrogates were included inside the pore or NTD region. After equilibration (detailed in Materials and Methods) the dodecameric structure was stable in all simulations, as indicated by the small root mean square deviation (RMSD, Extended Data Fig. 11 a,d).

The pore opening observed in our cryo-EM structures has a solvent-accessible radius of ∼3 Å (Figure 2b). This makes it the most narrow pore opening observed for a connexin channel to date (a comparison of the pore openings in the cryo-EM structures of connexin channels is shown in Extended Data Fig. 12). However, the average solvent-accessible radius of the pore during molecular dynamics was ∼6 Å (Figure 5c); note that the effective hydrated radius of K^+^ and Cl^-^ is ∼3.3 Å and ∼3.6 Å, respectively. The NTD regions of Cx43 are flexible (Extended Data Fig. 13) and can move laterally and vertically (Movie 1), allowing the passage of ions. The charge distribution inside Cx43 resembles that of the other GJCs ^18,19,37^ with a positive electrostatic potential in the NTD region of both HCs within a channel and a negative/neutral region near the HC interface (Fig. 5c). When no transmembrane potential is applied (0 mV, Fig. 5b), Cl^-^ ions accumulate in the NTD regions of Cx43 GJC (anion density peaks in the NTD region) as previously observed for Cx31.3 ^37^. At 0 mV, there is an entry barrier of ∼3.38 kcal/mol for cations (K^+^) in the NTD region of both HCs (Figure 5d) which is slightly higher than previously reported PMFs values for other homomeric GJCs (Cx46/50) ^18^. Conversely, anions can permeate the NTD region barrierless (Figure 5d), but they face a higher barrier ∼3.84 kcal/mol below this region where the electrostatic potential of the channel is negative, as observed for other GJC ^18^. Because the NTD domains are flexible, but they do not fully fold inwards, most ions enter and exit the same HC (Extended Data Table 2). Application of a transjunctional voltage lowers the barrier on one HC increasing the number of permeation events. Very few full transjunctional permeation events were observed during the simulations with any applied voltage.

**Figure 5.**
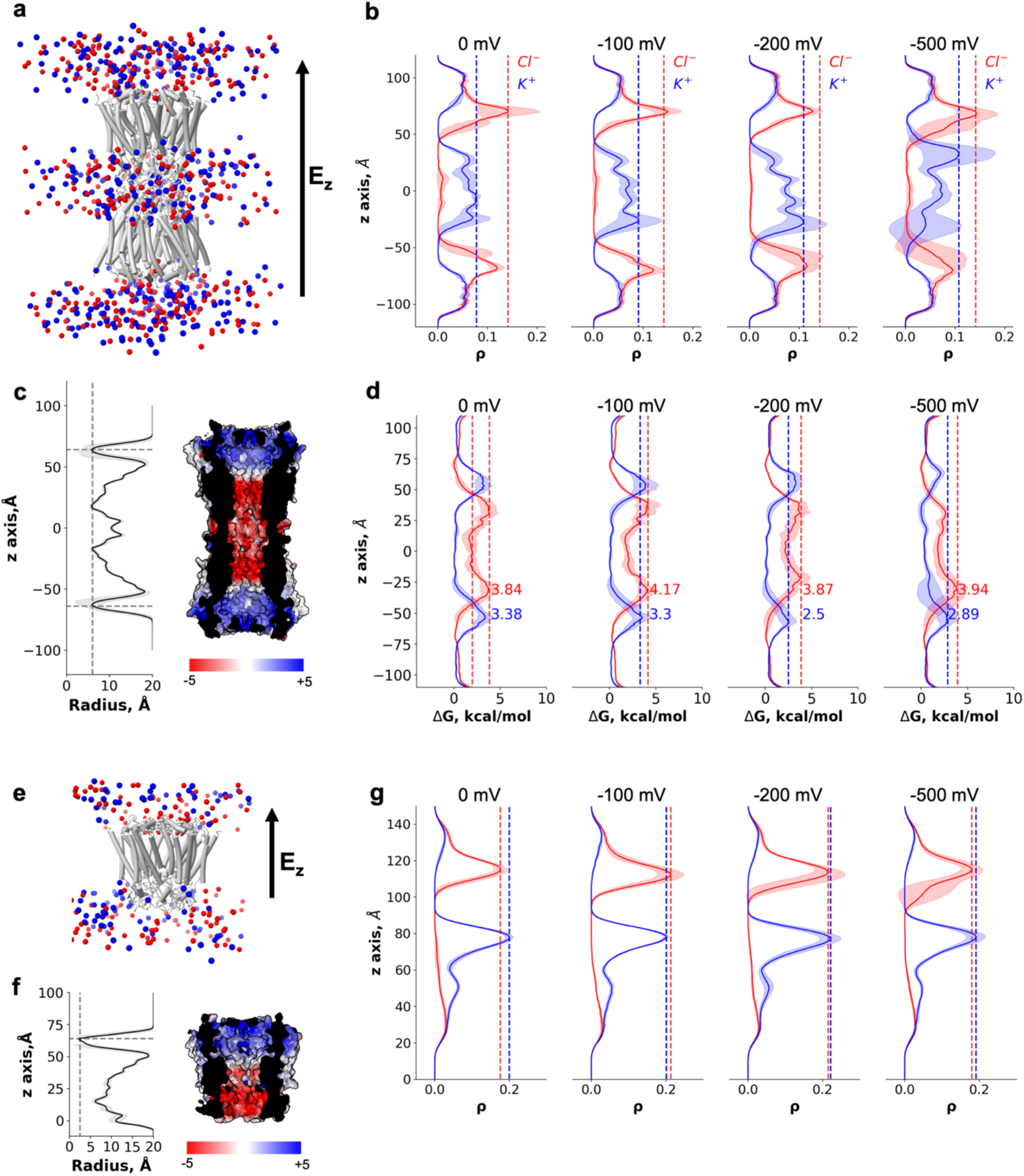
Molecular dynamics simulations of the Cx43 GJC and HC. **a**, Cartoon representation of the double-bilayer system including the GJC (cartoon) and ions (vdw spheres). Lipidic membrane and water residues have been removed for clarity. The direction of the applied constant electric field, E_z_, is indicated with an arrow. **b**, Ion density (ρ) profiles (average and fluctuations) along the diffusion axis of the GJC coloured in red for anions and blue for cations, for the simulated applied transjunctional voltages. **c**, The solvent-accessible radius (and fluctuations during MD) along the diffusion axis of the GJC. The dotted lines correspond to the minimum radius and the position of the NTD regions. The surface representation of Cx43 is coloured according to the calculated electrostatic potential; the slab view shows the properties of the pore. **d**, Average free energy experienced (at least two simulation replicates per panel) by the K^+^ (blue) and Cl^-^ (red) while permeating the pore with different applied transmembrane potential (from right to left: 0 mV, -100 mV, -200 mV and - 500 mV). The dotted lines correspond to the maximum free energy barrier for anions (red) and cations (blue). **e**, Cartoon representation of the lipid-bound system including the HC (cartoon) and ions (vdw spheres). Lipidic membrane and water residues have been removed for clarity. The direction of the applied constant electric field, E_z_, is indicated with an arrow. **f**, Same as c, for the Cx43 HC. **g**, Ion density (ρ) profiles (average and fluctuations) along the diffusion axis of the HC coloured in red for anions and blue for cations, for the simulated applied transmembrane voltages.

As mentioned before, the GJC did not fully open in any of our simulations, i.e., a conformation with all NTDs within the channel symmetrically moving to open the pore was not established at any point. To illustrate this effect, we calculated the root-mean-square deviation for each NTD domain in all simulations (Extended Data Fig. 13). Only at high applied transjunctional voltages only one NTD domain adopted a fully open folded inward conformation (Extended Data Fig. 13).

The MD simulations in the absence of a bound ligand reveal an intermediate, metastable state in which ions can permeate from both HCs with similar free-energy barriers and full transjunctional permeations are very rare events. The GJC selectivity and conductance properties are modulated by complex mechanisms involving both the steric aperture and the unique pattern of electrostatic features. In our case, the electrostatic properties of the channel are similar to those published for the Cx46/50 channels ^18^, but the steric contribution of the NTD region is significantly higher, increasing the barrier for K^+^, resulting in smaller ΔΔG differences than those observed for the open channel (Cx46/50). Other MD studies have shown similar energy barriers for ions ^18^, leading to entry events but very rare full translocations on the molecular dynamics timescale. Therefore, in the absence of the ligand the channel may exhibit a low residual conductance, as described experimentally ^38^.

Despite not knowing the precise identity of the lipid-N, we performed nine MD simulations of the Cx43 HC with a DHEA molecule as a lipid-N surrogate bound between the adjacent chains (Fig. 5e, Extended Data Fig. 14a,b, Movie 2), to understand whether the channel is closed in the presence of the lipid molecule. As expected, even when applying high transmembrane voltage the ion density for both species, K^+^ and Cl^-^, in the NTD region remained zero and no permeation events were observed (Fig. 5e-g, Extended Data Fig. 14c). The average pore radius during the simulations was consistent with that observed in the cryo-EM structure (Fig. 5f). Hence, the presence of the bound lipid-N molecule seals the channel by rigidifying the NTD domains.

## Discussion

The structures of Cx43 we have obtained by cryo-EM reveal a novel conformation of a connexin channel (both GJC and HC) featuring a closed molecular gate. The state is distinct from the previously observed structures of connexin GJCs and HCs, adding an important missing component to our understanding of connexin-mediated intercellular communication. With these structures in mind, we can now propose the existence of several structurally defined gating substates of the connexin channels: (i) a fully open gate, (ii) a semi-permeable central gate, and (iii) a closed gate (Fig. 6). The gating mechanisms of connexin HCs and GJC are complex and involve multiple components. The available evidence indicates that connexin channels are gated by membrane potential (V_m_), by transjunctional potential (V_j_), as well as by a range of chemical agents such as pH, Ca^2+^ and organic molecules ^38^. For example, a ball-and-chain model has been proposed for Cx26, whereby at low pH the NTDs extend deep into the pore and occlude the substrate translocation pathway ^39^. This conformation is distinct from the one captured by our cryo-EM reconstructions, and it is possible that under certain conditions the Cx43 NTD may transition from the observed conformation to a low pH Cx26-like state. It is worth mentioning that for the Cx43 HCs the open probability has been shown to be very low ^40^, and thus a closed conformation such as the one described here may represent the predominant state of the channel.

**Figure 6.**
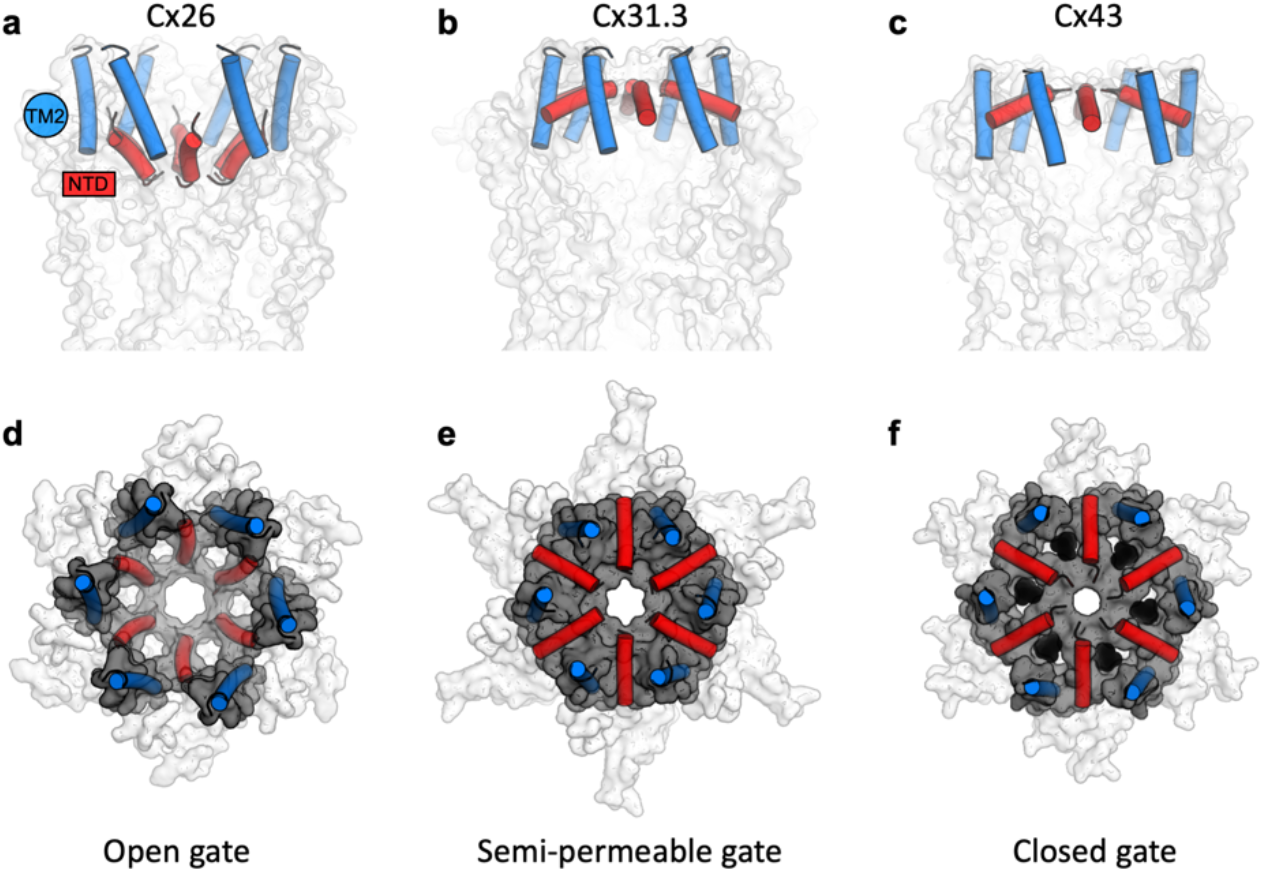
A structure-based view of connexin gating states. **a**, The side view of a fully open gate, as observed in Cx26 GJC (PDB ID: 2zw3); the gate-forming regions of the protein, NTD (red) and TM2 (blue), are shown as cylinders. Similar arrangement of the gating elements has been observed in Cx46/Cx50. This conformation of the gate is permeable to a wide range of substrates, including small molecules and ions. **b**, The semi-permeable central gate is featured in the structure of Cx31.3 HC (PDB ID: 6l3t). In this conformation the gate is likely selective for ions (Cx31.3 has been shown to have a preference for anions), based on the dimensions of the gate. **c**, The putative closed state is featured in the Cx43 structures (here, the Cx43 in digitonin is shown as an example). **d-f**, Same as a-c, viewed from the cytosolic side of the channel. The gray surface corresponds to the NTD and TM2 regions; the white surface corresponds to the rest of the protein; lipid-N molecules in f are shown as black spheres, using a model of DHEA manually placed into the six NTD lipid site regions for illustration purposes (the exact identity of the lipid-N molecule is unknown).

The cytosolic surface of the gate in our structures is positively charged, indicating that specific interaction with anions may be relevant to some of the gating events. As Cx43 is capable of cation and anion permeation ^41^, the presence of the positively charged surface in this region does not indicate ion selectivity. The differences in the electrostatic properties of the gates and the pore regions in different connexin channels may reflect the differences in their selectivity for ions and small molecules (Extended Data Fig. 12). However, it is also likely that the unstructured regions in Cx43 and other connexins, including the intracellular loop and the C-terminal domain, contribute to the electrostatic potential, substrate selectivity and regulation of these channels. These domains are known to play important roles in connexin channel regulation ^42^. However, as these regions are unresolved in any of the available 3D reconstructions of connexin channels, the structural basis of their action remains to be determined.

The observation of lipid-like molecules bound inside the pore is intriguing. The use of detergents and lipids is currently a prerequisite for structural analysis of membrane proteins, and it is possible that under solubilization conditions the abundance of added detergent and lipid (CHS) forces the protein to interact with these molecules. Nevertheless, these interactions may correspond to the naturally occurring interactions with the endogenous lipids present in the lipid bilayer of the cell. The presence of such molecules could have important implications for HC or GJC assembly, substrate permeation and molecular gating. It remains to be determined whether Cx43 or other connexin channels harbour intra-pore lipids in the cellular membranes *in vivo*.

Distinct gating modes have been shown in the literature for both Cx43 HCs and GJCs, with fast transitions from an open state to a state of residual conductance, and slow transitions from an open to a fully closed state ^38^. The structures of Cx43 described here may represent the closed states, based on the cryo-EM- and MD simulation-based evidence. The closed state could be stabilized by a yet unknown agent that occupies the NTD lipid site formed by the arrangement of the NTDs and TM2 regions observed in our structures. It is worth noting that the majority of Cx43 channels in a GJ plaque are closed, with only a small fraction of the channels active ^43^. Detailed investigations of Cx43 and other connexin channels will be required to determine whether the gate conformation observed here is a common feature among different GJCs and HCs, and to pinpoint the physiological circumstances that establish this gating conformation. Finally, capturing the structures of Cx43 channels in distinct conformations under a wide range of physiologically relevant conditions will be required to establish its mechanism of action, gating and regulation.

## Acknowledgements

We thank Emiliya Poghossian (EM Facility, PSI) and Miroslav Peterek (ScopeM, ETH Zurich) for expert support in cryo-EM data collection. We also thank Spencer Bliven and Marc Caubet-Serrabou (PSI) for the support in high performance computing. The work was supported by a grant from Horten Foundation, and by the Swiss National Science Foundation grant 184951 (VMK). SAG acknowledges support from an AGAUR Beatriu de Pinós MSCA-COFUND Fellowship (project 2020-BP-00177). FLG and SAG thank the CSCS and PRACE for supercomputing resources (projects pr126 and s1107).

## Competing interests

Authors declare that they have no competing interests.

## Data and materials availability

The atomic coordinates and structure factors have been deposited in the Protein Data Bank (7Z1T, 7Z22, 7Z23); the density maps have been deposited in the Electron Microscopy Data Bank (EMD-14452, EMD-14455, EMD-14456). The mass spectrometry data have been deposited at ProteomeXChange via PRIDE (PXD033824). The MD trajectories will be uploaded to Zenodo. All other data are available in the main text or the supplementary materials.

## Author contributions

C.Q. planned and performed the experiments, analyzed the data, wrote the manuscript, S.A.G. planned, performed and analyzed the simulations, co-wrote the manuscript, P.L. performed the experiments, co-wrote the manuscript, A.O. performed the experiments, analyzed the data, D.L.P. performed the experiments, analyzed the data, E.B. performed the experiments, analyzed the data, D.S. performed the experiments, analysed the data, co-wrote the manuscript, P.P. provided the reagents, equipment and expertise for mass spectrometry experiments, N.Z. provided the reagents, equipment and expertise for lipidomics analysis, M.B. planned the experiments, analyzed the data, co-wrote the manuscript, F.L.G. planned and analyzed the simulations, co-wrote the manuscript, V.M.K. planned and performed the experiments, analyzed the data, wrote the manuscript.

## Notes

### Competing Interest Statement

The authors have declared no competing interest.

### Summary of Updates

S.A.G. affiliation updated and funding information added to Acknowledgements. Fig. 2 labels fixed. Fig. 5f updated with a pore radius calculation for the Cx43 HC simulation. The gating region is now referred to as "putative".

