## Supplementary Information for "Structure of the connexin-43 gap junction channel in a putative closed state"

### Materials and Methods

**HeLa cell culture and immunostaining.** A clone of HeLa cells (DH clone, Sigma-Aldrich) that is essentially devoid of connexins was used for immunostaining and electrophysiology. Dual patch-clamp experiments (parental cells of Extended Data Fig. 1) confirmed that GJs are not present in these cells. HeLa cells were cultured in DMEM medium (Gibco) supplemented with 10% FBS (Gibco) and 1% PenStrep at 37 °C in a humidified incubator with 5% $CO_2$. The day after plating cells onto 12 mm diameter glass coverslips, they were transfected with a Cx43-IRES-YFP plasmid carrying the coding region of human Cx43 using Lipofectamine 3000 (Thermo Fisher Scientific).

Forty-eight hours after transfection, the cells were briefly washed with PBS and fixed using ice cold 2% PFA in PBS for 15 minutes. The cells were then permeabilized and saturated in 2% BSA and 0.01% Tween 20 PBS solution for 1 hour at room temperature. Cells were then incubated with anti-Cx43 primary antibody (AB1727, Sigma-Aldrich, 1:100 dilution) dissolved in 1% BSA in PBS for 3 hours at room temperature. After 3 washes, a secondary antibody (Atto647N goat anti–rabbit IgG, 40839-1ML-F, Sigma-Aldrich,1:250 dilution) was applied for 1 hour at room temperature. After brief washing and incubating the cells with DAPI (Merck, 1:100 dilution), the coverslips were mounted onto glass slides using Mowiol (Sigma-Aldrich) mounting medium. Fluorescence images were acquired using a Zeiss LSM 900 confocal microscope and an oil–immersion objective (63x / NA 1.4).

**Dual patch-clamp electrophysiology.** Twenty-four hours after transfection, a glass coverslip with Cx43-IRES-YFP transfected HeLa cells was transferred to an experimental chamber at room temperature (22–24 °C) and mounted on the stage of an upright wide-field fluorescence microscope (Olympus BX51WI) with an infinity-corrected water immersion objective (40X, 0.8 NA, Olympus). We continuously perfused cells at 2 ml/min with an extracellular solution containing 150 mM NaCl, 10 mM HEPES, 5 mM KCl, 5 mM glucose, 1 mM $MgCl_2$, 2 mM $CaCl_2$ and 2 mM sodium pyruvate (pH 7.4, 311 mOsm). Cytosolic YFP fluorescence was excited by a 460 nm LED to identify the transfected cells and allow subsequent electrophysiological analysis. Patch-clamp recordings were performed using an Axon 700B amplifier (Molecular Device) with a dual headstage configuration capable of carrying out simultaneous measurements. Pipettes were filled with an intracellular solution containing 130 mM KAsp, 10 mM NaCl, 10 mM HEPES, 10 mM KCl, 1 mM $MgCl_2$ and 50 μM BAPTA (pH 7.2, 305 mOsm) filtered through a 0.22 μm pore size membrane (Millipore). DC resistance of patch pipettes in the bath ranged from 6 to 8 MOhm. Once the whole-cell configuration was achieved, both cells were voltage-clamped at their common resting potential (around -20 mV). The junctional current $I_j$ was measured in cell 1 by applying voltage steps $V_j = +10$ mV to cell 2. The corresponding junctional conductance was computed as $g_j = I_j/V_j$ , without considering the potential drop $V_a$ due to pipette access resistance $R_a$, whose correction would be a source of numerical artifacts due to the large junctional currents involved. At the end of almost all experiments, we applied $CO_2$ to the bath to prove that junctional currents occurred through Cx43 GJ channels. Only cell pairs that showed complete uncoupling by the $CO_2$ were retained for the analysis. Electrophysiological data were acquired by pClamp software (version 10.4, Molecular Device) and analyzed with a software we developed in Matlab (The MathWorks, Inc.).

**Connexin 43 expression.** Human connexin 43 (Cx43, Uniprot ID P17302) with a C-terminal 3C-EYFP-twinStrep tag, cloned into pACMV plasmid, was used for protein expression in HEK293F cells. The cells were grown in Dulbecco's Modified Eagle's medium (DMEM) supplemented with 10% FBS and penicillin/streptomycin (PenStrep), in 15 cm plates at 37°C. Prior to transfection, the medium was exchanged to DMEM supplemented with 2% FBS and penicillin/streptomycin. The transfection mixture was prepared by mixing the expression vector and branched polyethyleneimine (PEI; Sigma Aldrich) at a ratio of 1:2 (w/w). After 5 min incubation, the transfection mixture was added to the cells. After 48 h the cells were collected using a cell scraper, washed with PBS by centrifugation and frozen at -80 °C until the day of the experiment.

**Connexin 43 purification.** The cell pellets were resuspended in buffer A (25 mM Tris-HCl, pH 8.0, 150 mM NaCl) supplemented with protease inhibitors (1 mM benzamidine, 1 µg/ml leupeptin, 1 µg/ml aprotinin, 1 µg/ml pepstatin, 1 µg/ml trypsin inhibitor and 1 mM PMSF), and lysed using sonication with a Vibra-Cell Sonicator (using 0.5 s pulses at 35% amplitude for 4 min). The cell membranes were collected by ultracentrifugation (Beckman Coulter Ti45 rotor, 35000 rpm). The membranes were resuspended using buffer A with protease inhibitors (1 mM benzamidine, 1 µg/ml leupeptin, 1 µg/ml aprotinin, 1 µg/ml pepstatin, 1 µg/ml trypsin inhibitor and 1 mM PMSF) and solubilized using a mixture of 1% DDM and 0.2% cholesteryl hemisuccinate (CHS) for 1 hour at 4 °C with rotation. After another round of ultracentrifugation, the supernatant was collected and mixed with 1 ml of CNBr-activated Sepharose coupled with anti-GFP nanobody [1]. Following a 30 min incubation at 4 °C, the resin was collected and washed with 40 column volumes of buffer B (25 mM Tris-HCl, pH 8.0, 150 mM NaCl, 0.1% digitonin). The protein was eluted overnight by HRV 3C protease cleavage (200 µg) at 4 °C. The eluted protein was concentrated using a 100 kDa cutoff Amicon concentrator and further purified by size exclusion chromatography (SEC) using Superose 6 Increase 10/300 GL column pre-equilibrated with buffer B. The peak fractions, corresponding to the protein of interest, were collected, concentrated, and used for cryo-EM grid preparation.

**Connexin 43 reconstitution in lipid nanodiscs.** For nanodisc reconstitution, Cx43 protein after GFP nanobody purification was concentrated to about 300 µl. The reconstitution system was Cx43 : MSP2N2 : POPC = 6 : 4 : 300. POPC was first mixed with Cx43 for 30 min at room temperature, then MSP2N2 was added for further incubation at room temperature for 30 min. Bio-beads (Bio-rad, 0.5 mg) were added to the reconstitution system. The sample was transferred to 4 °C and rotated for overnight to remove the detergent. The protein was centrifuged and loaded onto Superose 6 Increase 10/300 GL column pre-equilibrated with buffer A. The peak fractions, corresponding to Cx43 nanodisc, were collected, concentrated, and used for cryo-EM sample preparation.

**Western blot.** For western blot analysis, HEK293F cells were seeded in 6-well plates at a density of 500 000 cells/well in 10% FCS PSA DMEM and placed at 37 °C and 5 % CO2 over night to allow cells to adhere. The next day the medium was exchanged to 2 % FCS PSA DMEM and the cells transfected using branched PEI, with DNA to PEI at 1:2 ratio. The cells were placed at 37 °C and 5 % CO2 for 36 h expression.

The cells were washed twice with PBS and resuspended in 500 µl of PBS. The cells were pelleted by centrifugation at 1000 g for 15 min at 4 °C and resuspended in 300 µl of 25 mM Tris-HCl pH 8.0, 150 mM NaCl, supplemented with protease inhibitors and DNase I. The cells were sonicated by a single pulse (0.5 s on, 0.5 s off) at 35 % amplitude and 4X SDS PAGE loading buffer was added to the sample. Equal volume of sample was loaded on 4-20 % MiniPROTEAN TGX gradient SDS PAGE gel (Biorad) and the gels were ran at 150 V for 45 min. The transfer to nitrocellulose membrane was done in Trans-Blot SD cell (Biorad) using 50 mA per gel for 1 h in 48 mM Tris, 39 mM glycine, 20 % methanol and 0.04 % SDS. The membrane was blocked in 5 % BSA TBST (0.1 % Tween20) for 1 h, and incubated with rabbit anti-Cx43 antibody (ab217676, Abcam) diluted at 1:1000 in TBST (0.1 % Tween20) at 4 °C over night at constant rotation. The membranes were washed three times with TBST (0.1 % Tween20) and stained with goat anti-rabbit HRP conjugated secondary antibody (ab6721, Abcam) at 1:10000 dilution in TBST (0.1 % Tween20). The membranes were washed again three times with TBST (0.1 % Tween20), the signal developed using SuperSignal West Pico

PLUS Chemiluminescent Substrate and imaged using Amersham Imager 600 using automatic exposure.

**Mass spectrometry-based protein characterization**

*Protein digestion.* A sample of purified protein (25 µg) was digested using ProtiFi S-Trap™ micro spin columns according to the manufacturer's protocol. After protein digestion the sample was dried in a vacuum concentrator and resuspended in 725 µL 5% acetonitrile (ACN), 0.1% formic acid (FA). Part of the sample was transferred to an HPLC vial and further diluted 1:2 prior to analysis.

*LC-MS/MS data acquisition.* 1 µL of sample was analyzed on an Orbitrap Eclipse™ Tribrid™ Mass Spectrometer (Thermo Fisher) equipped with a nanoelectrospray source, connected to a nano-flow LC system (Easy-nLC 1200, Thermo Fisher). Peptides were separated on a 40 cm x 0.75 µm (inner diameter) column packed in-house with 1.9 µm C18 beads at a flow rate of 300 nL/min and a 60 min linear gradient from 3-30% II (Eluent I: 0.1% FA, Eluent II: 95% ACN, 0.1% FA). The column was heated to 50 °C. The sample was measured three times in data-independent acquisition (DIA) mode with 41 variable width DIA windows with a 1 m/z overlap. For survey MS1 spectra the measured mass range was 350-1400 m/z at a resolution of 120,000 with 200% normalized AGC target or 100 ms maximum injection time. Survey MS1 spectra were acquired in between complete DIA isolation window sets. MS2 spectra covered a mass range of 150-2000 m/z at a resolution of 30,000. HCD collision energy for MS2 spectra was set to 30% with 400% normalized AGC target or 54 ms maximum injection time.

*Peptide and protein identification.* Peptide and protein identification of DIA measurements was performed using Spectronaut™ software [2] (Biognosys, version 14.5) in directDIA™ mode. Default settings were used with minor adjustments. Minimum peptide length was set to 5 amino acids and Phospho (STY) was included as a variable modification. The results of the analysis are shown in Extended Data Figure 5.

**Lipidomics analysis of purified protein.** Cx43 was purified in DDM using GFP nanobody affinity chromatography with a buffer buffer 25 mM Tris-HCl, pH 8.0, 150 mM NaCl, 0.02% DDM. Non-transfected HEK 293F cells were used as a mock purification using the same purification method. The lipid was extracted by butanol-methanol single phase extraction. The extraction buffer was a mixture of $H_2O$: butanol: methanol at a volume ratio of 2:9:9. 100 µl protein sample (about 0.1 mg) was mixed with 900 µl extraction buffer and incubated at room temperature for 30 min with vortexing every 5 min. After incubation, the sample was centrifuged at 10000 rcf/min for 10 min. The supernatant was transferred to a glass vial and used for mass spectrometry analysis.

Liquid chromatography was done as described previously [3] with some modifications. The lipids were separated using C18 reverse phase chromatography. Vanquish LC pump (Thermo Scientific) was used with the following mobile phases; A) ACN:Water (6:4) with 10 mM ammonium acetate and 0.1% FA and B) Isopropanol:ACN (9:1) with 10 mM ammonium acetate and 0.1% FA. The Acquity BEH column (Waters) with the dimensions 100 mm * 2.1 mm * 1.7 µm (length * internal diameter * particle diameter) was used. The following gradient was used with a flow rate of 0.6 ml/min; The following gradient was used with a flow rate of 0.6 ml/min; 0.0-2.0 min (ramp 15-30% B), 2.0-2.5 min (ramp 30-48% B), 2.5-11 min (ramp 48-82% B),11-11.5 min (ramp 82-99% B), 11.5-12 min (isocratic 99% B), 12.0-12.1 min (ramp 99-15% B) and 12.1-15 min (isocratic 15% B).

The liquid chromatography was coupled to a hybrid quadrupole-orbitrap mass spectrometer (Q-Exactive HFx, Thermo Scientific). A full scan acquisition in positive ESI was used. A full scan was used, scanning from 200-2000 m/z at a resolution of 120000 and AGC Target 1e6 and a max injection time 200 ms, while data-dependent scans (top10) were acquired using normalized collision energies (NCE) of 20, 30, 50, a resolution of 15,000, and AGC target of 1e5. Identification of the lipids was achieved using three criteria: 1) High accuracy and resolution with an accuracy within m/z within 5 ppm shift form the predicted mass and a resolving power 70000 at 200 m/z; 2) Isotopic pattern fitting to expected isotopic distribution; 3) The fragmentation pattern matching lipidblast library [3] and mzcloud (ThermoScientific). Mass spectrometric data analysis was performed in Compound Discoverer 3.1 (ThermoScientific) for peak picking, annotation and matching to lipidblast.

**Cryo-EM sample preparation, data collection and processing.** The purified Cx43 protein in digitonin was concentrated to about 5 mg/ml. The Cx43 nanodisc protein was concentrated to about 1.5 mg/ml. A 3.5 µl aliquot of the protein was applied to a glow-discharged Quantifoil R1.2/1.3 200-mesh grid. The grid was blotted using Vitrobot Mark IV (Thermo Fisher Scientific) and plunge-frozen into liquid ethane. Three cryo-EM datasets were collected using a Titan Krios electron microscope equipped with a K3 direct electron detector and a GIF-Quantum energy filter (slit width of 20 eV) at ScopeM, ETH Zurich. The defocus range was set from -1.0 µm to -2.0 µm. The movies were collected in super-resolution mode using EPU 2.0, dose-fractioned in 40 frames. The exposure time for each micrograph was 0.8 s and the total dose was ~50 $e^-/Å^2$.

The images were assigned to distinct optics groups according to the EPU beam shift values using a script provided by Dr. Pavel Afanasyev (ETH Zurich; https://github.com/afanasyevp/afis). The movies were binned two-fold and motion-corrected using MotionCor2 [4], yielding aligned micrographs with a pixel size of 0.678 Å. CTF estimation was performed using Gctf [5]. Particle picking and 2D classification was performed in Relion 3.1 [6] (nanodisc sample) and Relion 4.0 [7] (detergent sample). After several rounds of 2D classification, the selected particles were used for 3D classification.

For detergent sample, the best 3D classes from the 3D classifications were merged and refined to 2.61 Å. After CTF refinement and particle polishing, the resolution was improved to 2.43 Å. An additional round of 3D classification without alignment was performed, the best class was selected for 3D refinement, yielding a 2.41 Å resolution refined map. Continued refinement with a detergent-free mask yielded a reconstruction at a resolution of 2.26Å. To further analyse the conformation viability, we performed particle symmetry expansion using *relion_particle_symmetry_expand* and subtracted the particles with a box size of 200 pixels. The subtracted particles were further classified using 3D classification in Relion with T=20. Two 3D classes showed high resolution features but were not substantially different from each other.

For Cx43 in nanodiscs, two datasets were processed separately for 3D classification. The best hemichannel classes were joined together for 3D refinement to 3.56 Å. Because of the preferential orientation, two rounds of 3D classification were performed. With a detergent-free mask, the final refinement reached 3.98 Å and yielded a less anisotropic map. This map was used for model building. For gap junction channel, the best classes from 3D classification were joined to 3D refinement to 3.25 Å. An additional round of 3D classification without alignment was performed. After CTF refinement, the resolution was improved to 2.95 Å. The pixel size was corrected during postprocessing to 0.654 Å. Local resolution maps were calculated by ResMap [8] implemented in Relion. Symmetry expansion and 3D classification was performed in Relion. The detailed steps of image processing are shown in Extended Data Figs. 3-4, 6-8 and Extended Data Table 1.

**Model building and refinement.** Model building was carried out manually in Coot [9]. Residues L106-G150, V236-I382 were not built because these regions were not resolved in the refined density map. The M1 residue was excluded based on the MS data, which indicate that the N-terminal peptides present in the sample lack the M1 (Extended Data Figure 5). The side chains of the bulky residues (Phe, Trp, Tyr and Arg) were used as a guide for model building, further aided by the available connexin structures. The model was refined using real_space_refine in Phenix [10]. The model was validated as previously described [11]. The atoms of the final model were randomly displaced by 0.5 Å using PDB tools implemented in Phenix. The perturbed models were refined in Phenix against the half map 1. The refined model was used to generate the FSC curve of the model versus half map 2. The geometry of the model was validated using MolProbity [12]. All figures were prepared in PyMol [13], Chimera [14] or Chimera X [15].

**Molecular dynamics simulations.** The CHARMM-GUI [16] server was used to build the initial simulation box comprising the full dodecameric gap junction Cx43 with one hemichannel embedded in a POPC bilayer and fully solvated using TIP3P water. Sidechains were protonated according to neutral conditions. Disulphide bonds, identified in the experimental structure, were enforced. Amino acids corresponding to the ICL (residues 106 to 151) and CTD (residues 236 to 382) domains were not modeled, as the experimental data describing these large disordered portions of the structure were inconclusive, and successful MD simulations have been carried out for other dodecameric gap junctions without including these structural features [17,18]. The second bilayer was added, replicating by symmetry the one embedded with CHARMM-GUI, the water placed inside the channel was kept and the rest of the system was solvated again using GROMACS [19] with 150 mM KCl. The final system was cleaned to remove unfavourable water positions. The topology was constructed using CHARMM36M [20] for the protein, POPC lipids, and TIP3P [21] water. The final system had 461767 atoms: 37320 protein atoms, 136144 POPC atoms, 287373 TIP3P atoms, 447 $K^+$ and 483 $Cl^-$ ions. The net charge of the gap junction is 36e-. We also built a system with counterions only.

The system was equilibrated using GROMACS2021 [19,22] starting from an initial minimization step with the protein and lipid molecules harmonically constrained during 5000 steps, followed by a 125 ps (dt=1fs) heating step at 303.15 K in the NVT ensemble. A second heating step of 125 ps (dt=1fs) was performed at 303.15 K in the NVT ensemble decreasing the lipid restraints. In the third step we decreased the lipid restraints and applied a Berendsen semi-isotropic barostat (1 atm) during 125 ps (dt=1fs) keeping the temperature at 303.15 K. In the fourth step (500 ps) we kept decreasing the restraints and increased the integration timestep to 2 fs, keeping a semi-isotropic barostat (1 atm) and thermostat at 303.15 K. We performed a final NPT step (30 ns) keeping the restraints on the $C_\alpha$ of the interface only. Production runs were performed with V-rescale thermostat [23] at (303.15 K) with periodic boundary conditions and particle mesh ewald (PME). Eighteen independent simulations (100 ns) of the GJC in the absence of ligands were started from the equilibrated structure.

The system with the DHEA molecule as a lipid-N surrogate modelled in the NTD lipid sites was equilibrated following the same protocol but the center of mass of the DHEA molecule was harmonically restrained during equilibration and production runs, same as the NTD, with a harmonic force constants of 1000 kJ mol-1 nm-2.

We performed constant electric field simulations [24] with different applied transjunctional voltages (-100 mV, -200 mV, -500 mV) where the applied electric field was calculated as $E_z=V_{applied}/L_z$ ($L_z$ is the simulation box length along the channel diffusion axis, z).

RMSD calculations were calculated using GROMACS2021. Interface distances were computed using PLUMED2.7 [25]. All MD plots were made with the seaborn library for python. The cartoon representations included in the panels of some figures and movies were created using ChimeraX [26]. For every trajectory, we computed the density of ions within a cylinder of radius 10 Å using an in-house code that provides the average (in time) number of ions in a 3D grid, spaced every 1 Å. For all grids, we computed the integral along the Cx43 diffusion axis (z, axis). We then computed the K+/Cl- free energy profiles (Figure 5) along the diffusion axis of the Cx43 channel as $\Delta G=-RT\ln(\bar{\rho})$ and its error as $\sigma\_\Delta G=|RT\ \sigma\_\rho/\rho|$.

Using the same code (described above for the ion density calculations) for the protein-only production trajectory, we computed the solvent-accessible area and fluctuations by counting the empty grid points.

**Cell culture for light microscopy experiments.** HEK293F cells were seeded at a density of 30 000 cells per well in poly-L-lysine pre-coated ibidiTreat µ-slide microscopy chambers for protein membrane colocalization studies in Dulbecco's Modified Eagle Medium (DMEM) supplemented with 10 % FBS and PenStrep. The cells were left over night at 37 °C and 5 % $CO_2$ and transfected the next day. They were transfected using branched PEI with DNA to PEI in 1:2 ratio (w/w). For plasma membrane colocalization studies, 0.46 µg of pAYST-Cx43 (Cx43 with a YFP tag) was used. DNA and PEI dilutions were prepared separately in DMEM medium, mixed and incubated for 5 min before added to the wells or dishes in a drop-wise manner. The cells were placed in the incubator at 37 °C and 5 % $CO_2$ for 48 h.

*Protein membrane colocalization studies.* The cells expressing Cx43-YFP were stained with Hoechst 33342 (50 µg/ml) and Vybrant CM-DiI (1:100 dilution) in 1 X PBS for 15 min at 37 °C and 5 % $CO_2$. They were washed two times with 1 X PBS and imaged in FluoroBright DMEM supplemented with 1 % FBS and PenStrep.

**Confocal light microscopy.** For all purposes, the cells were imaged using Leica Stellaris 5 confocal microscope equipped with a HyD detector, using LAS X (4.2.1.23819 – build 23180) acquisition software, frame sequential data acquisition scheme using 405 nm, 488 nm, 561 nm laser lines for Hoechst 33342, EYFP, CM-DiI, respectively. The images were collected with 20 X air objective (NA 0.75) and a pinhole size of 38.3 µm.

**Confocal light microscopy image analysis**

Co-localization studies were performed in Fiji [27] using the Coloc2 plug-in after background subtraction and 8-bit conversion of images. To confirm that the observed lines between two cells are Cx43-YFP GJCs located on the plasma membrane, the threshold Manders' co-localization coefficient [28] of YFP with CM-DiI was determined at those lines, defined as the regions of interest (ROI) ($n = 9$; mean ± SD = 0.57 ± 0.18). These coefficients were compared to coefficients of YFP in randomly selected cellular ROIs ($n = 9$; mean ± SD = 0.05 ± 0.09). The statistical significance was determined using unpaired t-test (P value < 0.0001, ****).

### Supplementary Figures

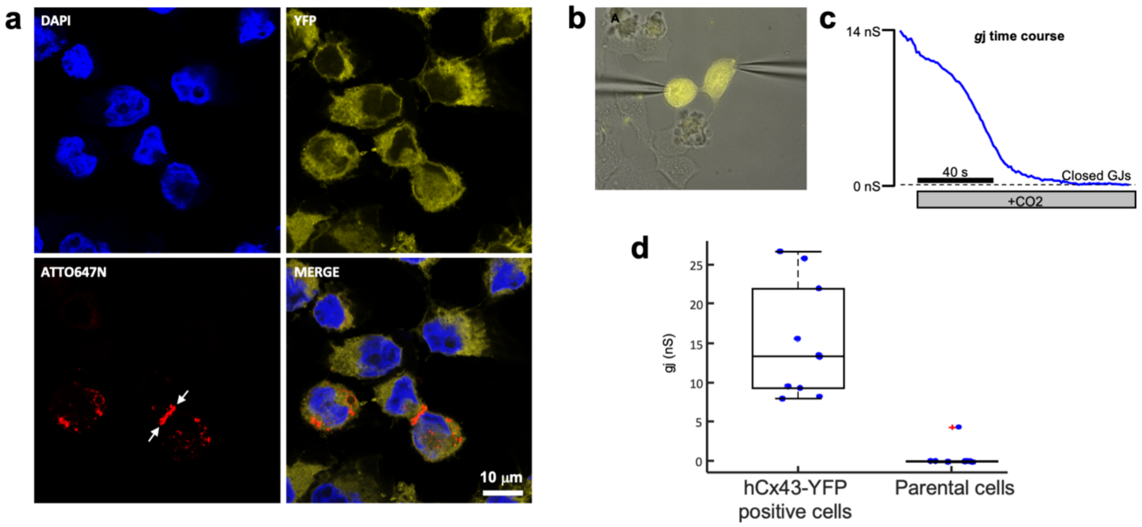


**Extended Data Figure 1. Localization and function of Cx43 gap junction channels (GJCs).** **a.** Immunostaining analysis of transfected HeLa cells expressing Cx43 (red) and cytosolic YFP (yellow). The red fluorescent image of anti-Cx43 ATTO647N-conjugated antibody indicates the presence of a large Cx43 gap junctional region (indicated by white arrows) between neighbouring cells. **b.** HeLa cell pair expressing high levels of cytosolic YFP selected to record Cx43 junctional current by dual patch-clamp. **c.** $g_j$ junctional conductance of the cell pair in b derived in the dual whole-cell configuration. Cell perfusion with saturated $CO_2$ extracellular solution was here used to verify that the junctional current was mediated by connexin channels. **d.** Box plot of $g_j$ measured in YFP-positive (n = 10) and parental (n = 10) HeLa cell pairs. Mean ± STD is 15.2 ± 7.2 nS and 0.5 ± 1.4 nS, respectively. Statistical analysis was performed using the Mann-Whitney $U$ test, yielding $p = 0.0002$.

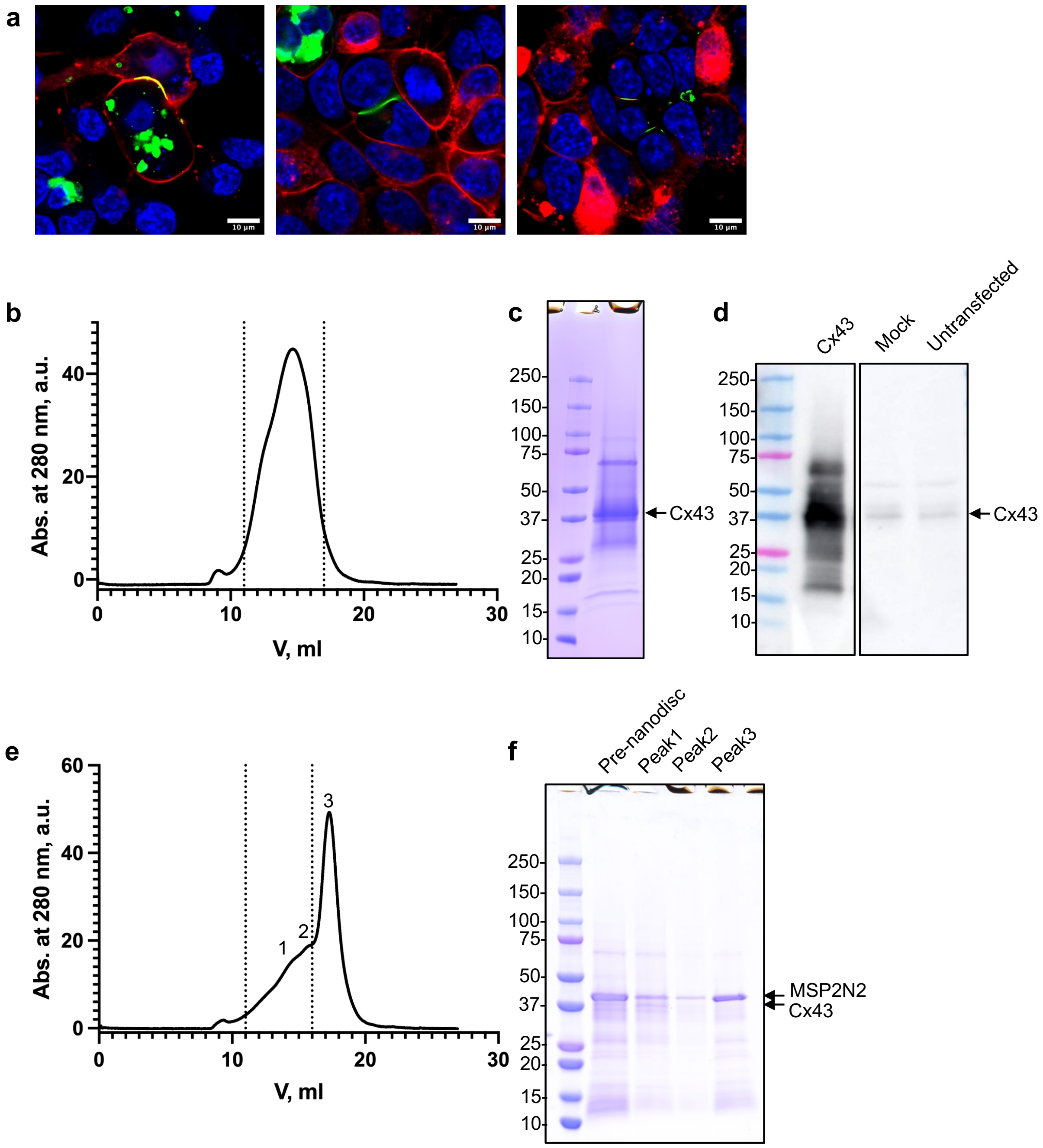


**Extended Data Figure 2. Expression and purification of Cx43. a.** Live cell microscopy images of HEK293F cells expressing Cx43; Cx43 was fused with a C-terminal YFP tag (green). The membrane was stained with Vybrant CM-Dil (red) and the nuclei were stained with Hoechst 33342 (blue). The gap junction was formed on the cell membrane between two cells. **b-c**, Size exclusion chromatography (SEC; b) and SDS-PAGE (c) of Cx43 purification. The dashed line indicates the fractions of the protein used for the experiments. **d**. Western blot analysis of lysates of cells expressing Cx43 (tag-free), using an anti-Cx43 antibody. For comparison, cells mock transfected with an empty plasmid and untransfected HEK293F cells are shown. The major band corresponding to the full-length Cx43 is similar to that in c. **e-f**. Same as b-c, for the MSP2N2 nanodisc-reconstituted Cx43 sample. The numbers correspond to the SEC fractions of the protein.

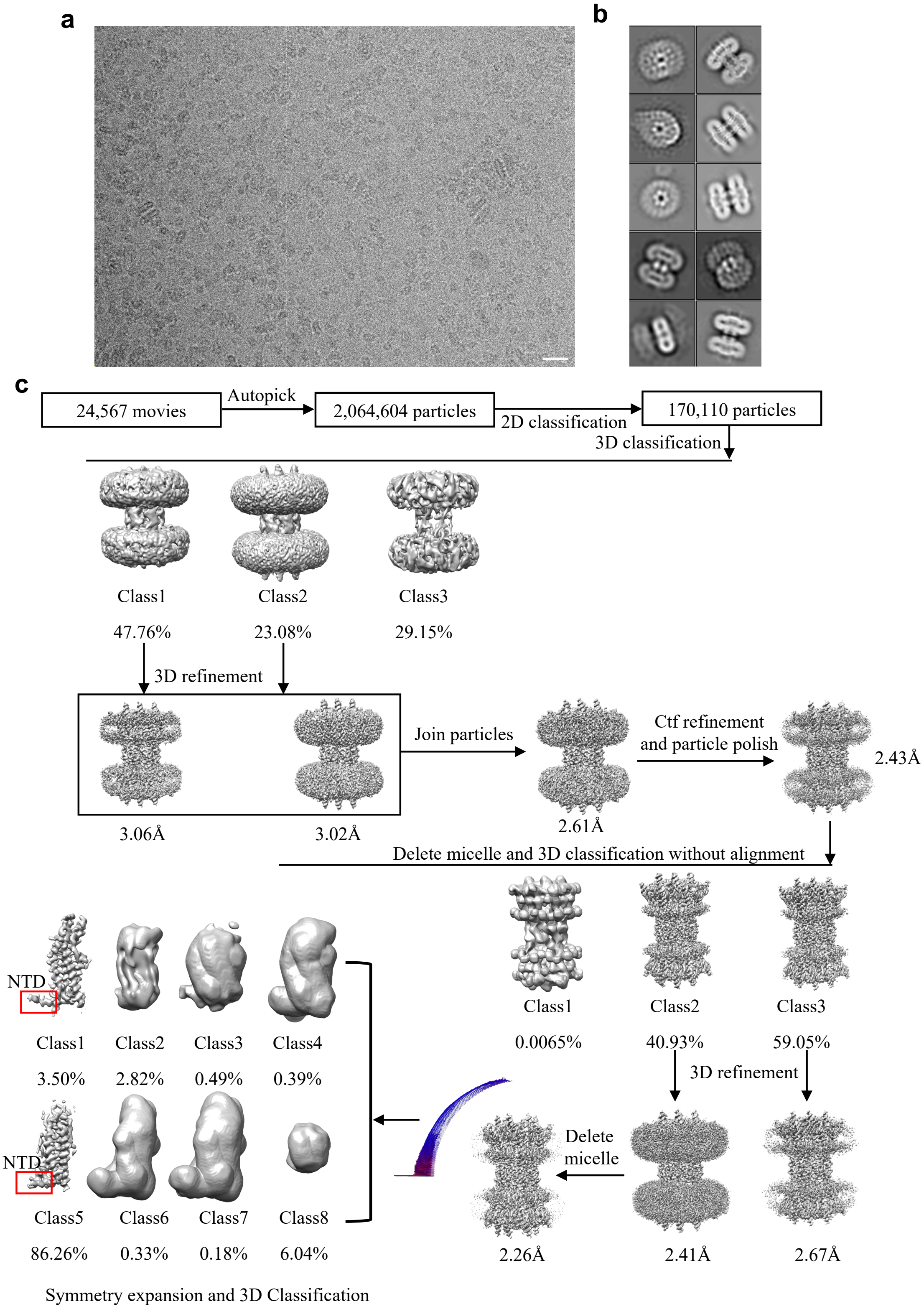


**Extended Data Figure 3. Cryo-EM image processing workflow of Cx43 in digitonin. a.** A representative aligned average micrograph of Cx43 sample; the scale bar corresponds to 20 nm. **b.** Representative 2D classes of Cx43; the box edge is 260Å. **c.** The overview of the Cx43 cryo-EM image process, including 2D, 3D classification and 3D refinement. Angular distribution histogram of the final reconstruction is represented at the bottom left. Symmetry expansion and 3D classification revealed that the majority of particles cluster into a single class (class 5); a minor class (class 1) showed a similar NTD arrangement.

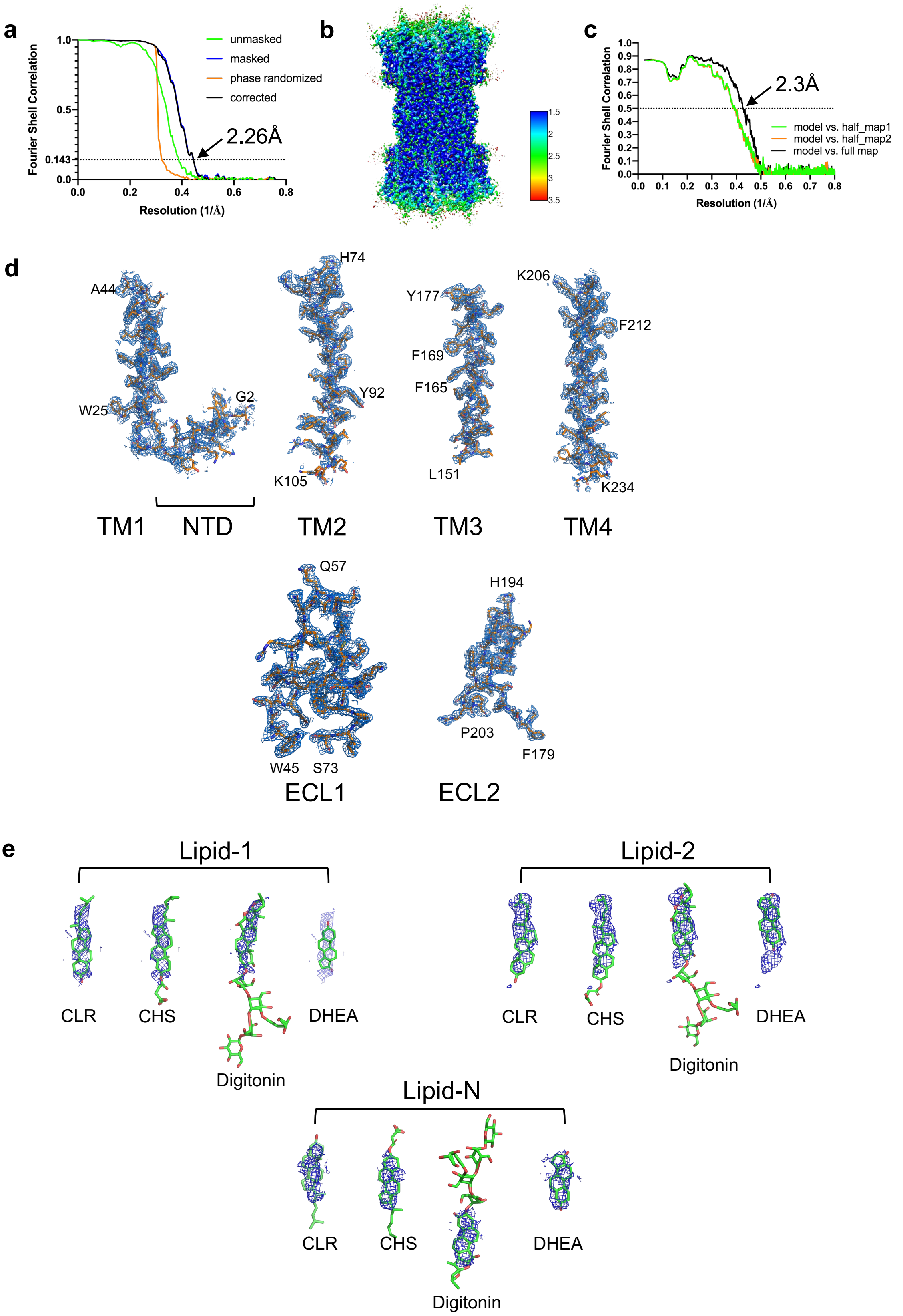


**Extended Data Figure 4. Fourier shell correlation (FSC) of the Cx43 reconstruction, local resolution and density map features. a.** Gold-standard FSC plot of Cx43 reconstruction. **b.** Local resolution map of Cx43, estimated using ResMap. **c.** Model to map FSC plot. **d.** Isolated

Cx43 density map regions of NTD/TM1, TM2, TM3, TM4, ECL1 and ECL2. **e.** A view of the Cx43 lipids densities fit with different possible ligands, CLR, CHS, Digitonin or DHEA.

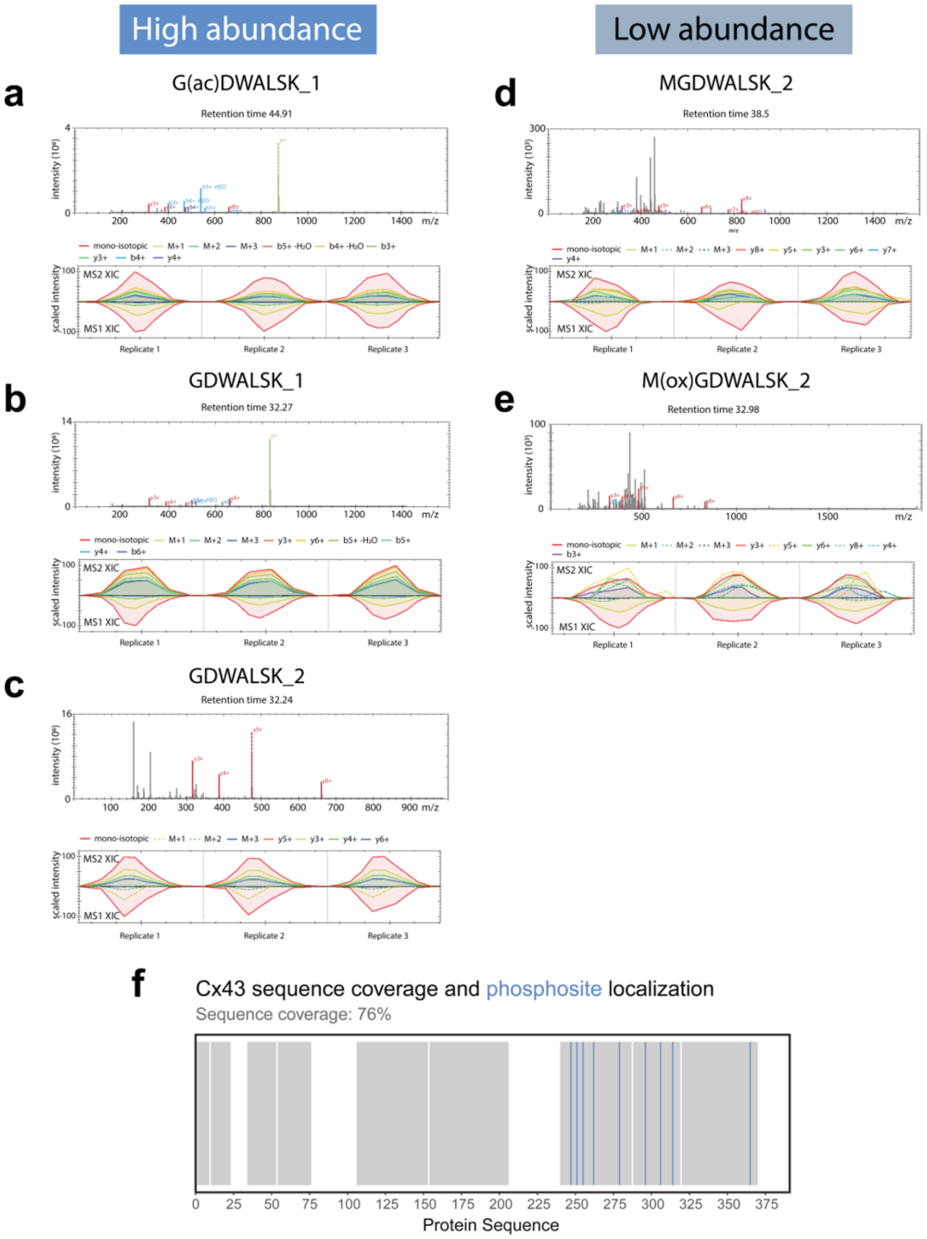


**Extended Data Figure 5. Mass spectrometric characterization of purified Cx43. a-e.** Fragment spectra for each of the identified peptides (upper panel) and extracted-ion chromatograms (XIC) for fragment and parent ions detected in MS2 and MS1 spectra (lower panel). **a-c.** An N-terminal peptide, $G_2DWSALGK_9$, was identified using LC-MS/MS data acquisition and analysis. A fraction of the peptide acetylated at the residue $G_2$ was also observed in the sample, suggesting a mixture of peptide species (c). **d-e.** The N-terminal peptides $M_1GDWSALGK_9$ and $M(ox)_1GDWSALGK_9$ were identified in low abundance, suggesting M1 was deleted during protein expression. **f.** Barcode plot showing the identified regions (grey) along the sequence of the purified Cx43 (residues 1-391) and the location of identified phosphosites (blue) with at least 75% confidence at positions 247, 251, 255, 262, 279, 296, 306, 314, 365.

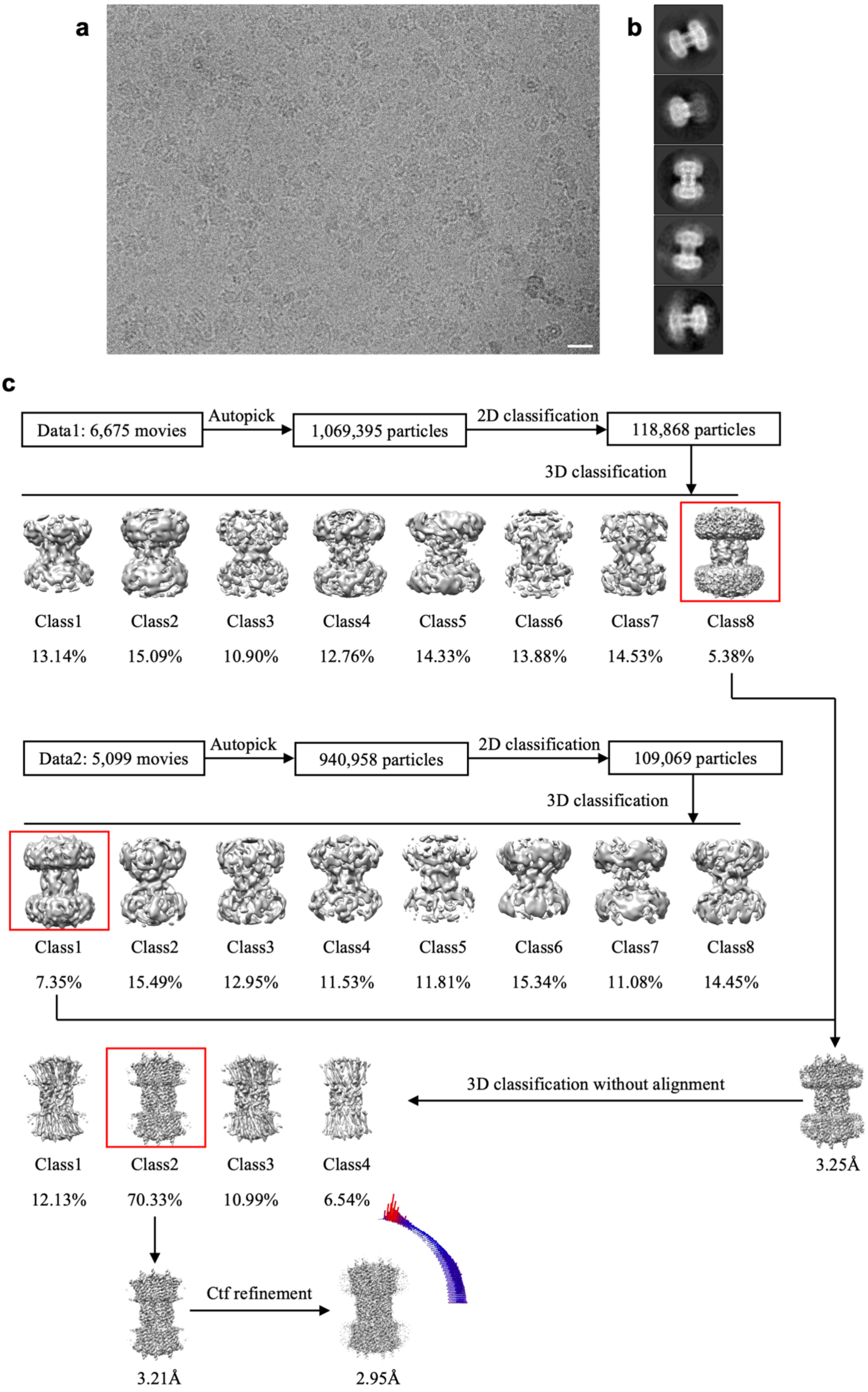


**Extended Data Figure 6. Cryo-EM image processing workflow for Cx43 gap junction channel (GJC) in nanodisc. a.** Representative 2D classes of Cx43 GJC in nanodisc sample; the box edge is 260.35Å. **b.** The overview of the Cx43 GJC in nanodisc cryo-EM image process, including 2D, 3D classification and 3D refinement. Angular distribution histogram of the final reconstruction is represented at the bottom right.

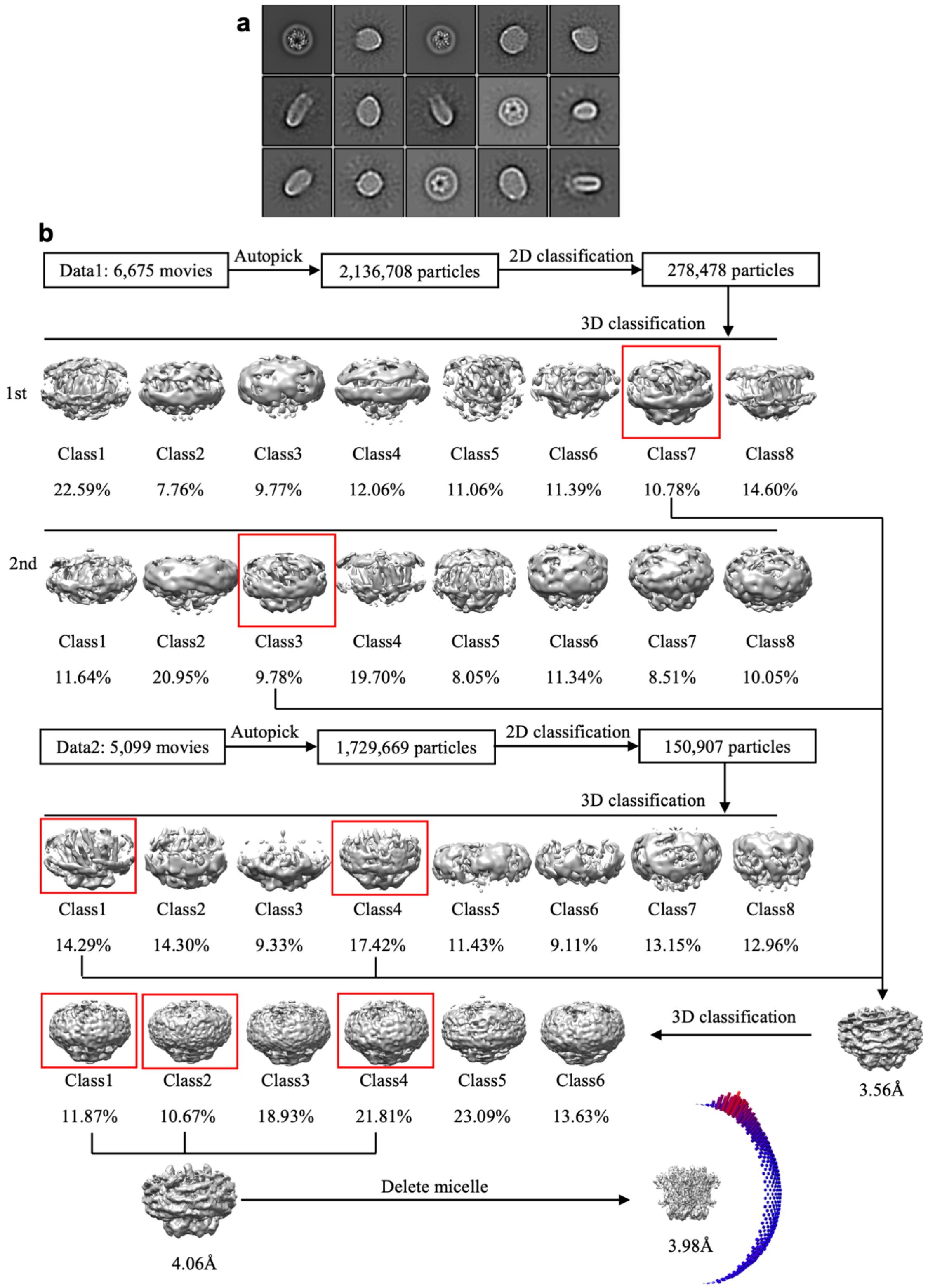


**Extended Data Figure 7. Cryo-EM image processing procedure for Cx43 hemichannel (HC) in nanodisc. a.** A representative motion corrected micrograph of Cx43 nanodisc sample; the scale bar corresponds to 20 nm. **b.** Representative 2D classes of Cx43 HC in nanodisc sample; the box edge is 260.35Å. **c.** The overview of the Cx43 HC in nanodisc cryo-EM image process, including 2D, 3D classification and 3D refinement. Angular distribution histogram of the final reconstruction is represented at the bottom right.

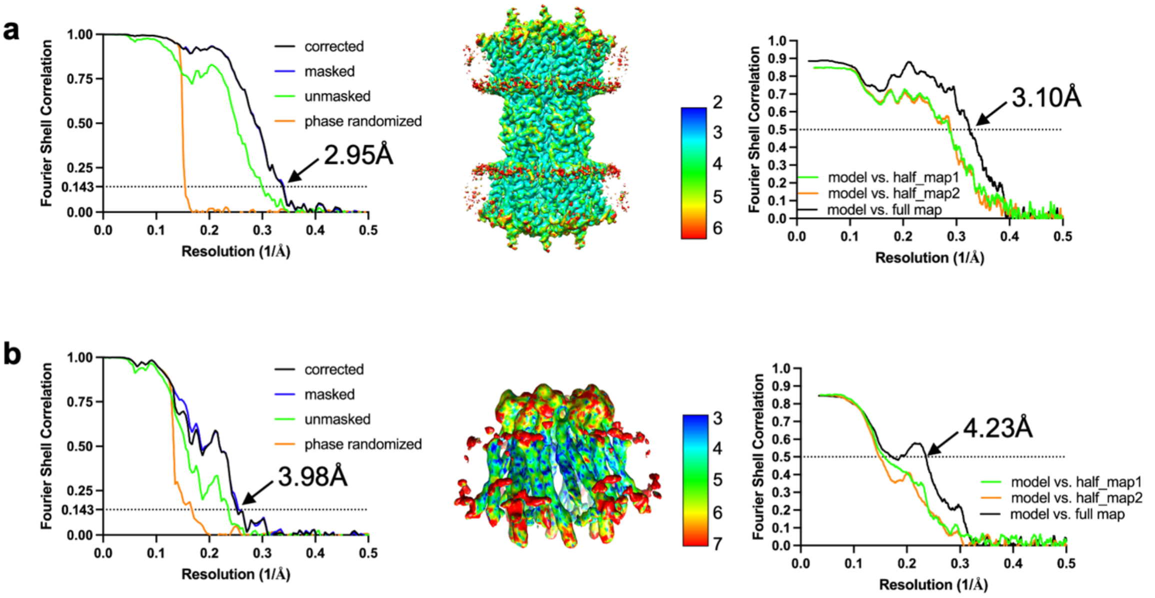


**Extended Data Figure 8. Fourier shell correlation (FSC) and local resolution maps of Cx43 in nanodiscs. a.** Gold-standard FSC plot, local resolution and model to map FSC of Cx43 GJC in nanodiscs. **b.** Same as a, for Cx43 HC in nanodiscs.

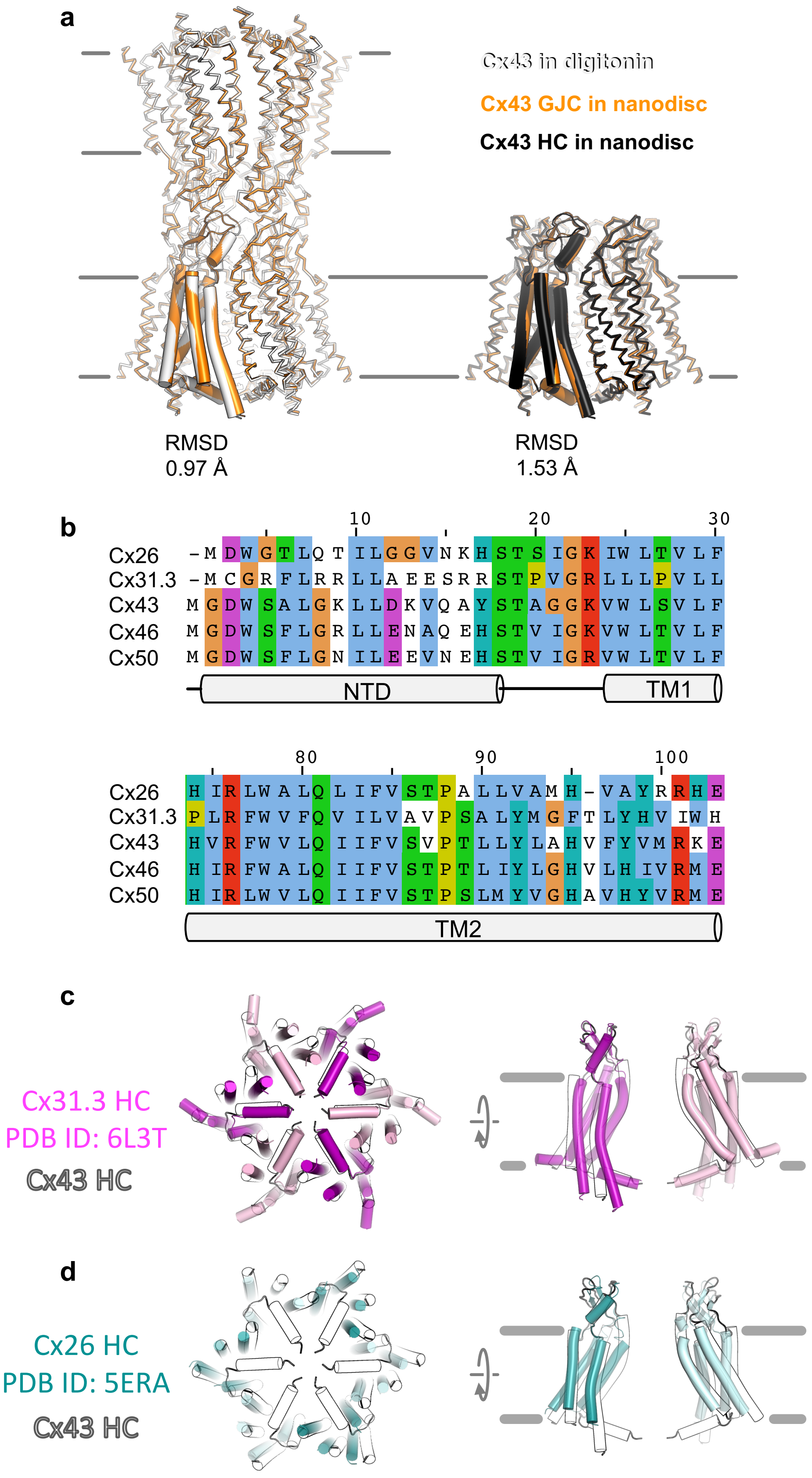


**Extended Data Figure 9. Comparison of the three cryo-EM structures of Cx43 and sequence alignment of NTD and TM2. a.** Structural alignment of Cx43 GJCs in digitonin and Cx43 in MSP2N2 nanodiscs (*left*), and for the free Cx43 HCs and the Cx43 HCs within the MSP2N2 nanodiscs (*right*). The RMSD between the corresponding structures in indicated in the panel. **b.** Sequence alignment of the two key structural elements: N-terminal domain (NTD) and TM2, which form the putative cytosolic gating region of Cx43. The sequences used here correspond to the proteins for which structures have been determined experimentally. **c.** A comparison of Cx43 HC (white contours) with the Cx31.3 HC structure (pink/magenta; PDB ID: 6L3T). **d.** Same as in c, with the Cx26 HC (cyan/teal, used as an HC approximation based on the Cx26 GJC structure; PDB ID: 5ERA).

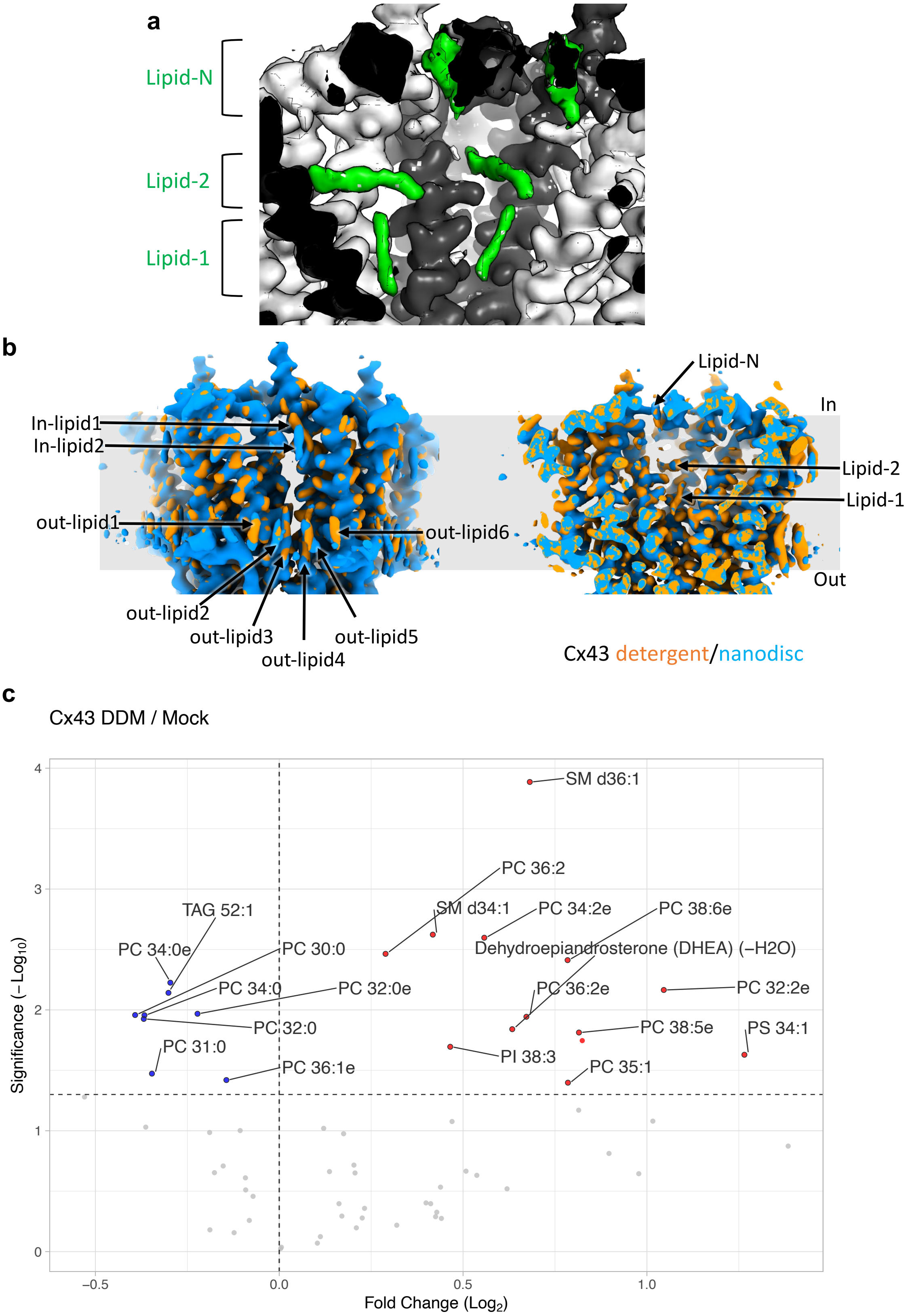


**Extended Data Figure 10. Comparison of intrapore lipid densities in detergent and in nanodiscs and lipidomic analysis of Cx43. a.** A view of the intrapore lipid densities in the density map of Cx43 GJC in nanodisc; the lipid densities are shown using the unsharpened refined density map, contoured at 4 σ. **b.** Comparison of the Cx43 density maps for the detergent and nanodisc samples reveals the conserved positions of the annular lipids (*left*) and intrapore lipids (*right*). **c.** Lipidomic mass spectroscopy analysis of Cx43 sample versus mock purification. Cx43 was purified in detergent (DDM) to avoid introducing any additional sterol molecules into the analysed sample. Mock-purified samples prepared from non-transfected HEK293 cells were used as a mock control. Lipidomic mass spectroscopy analysis was performed to identify lipid molecules co-purified with Cx43.

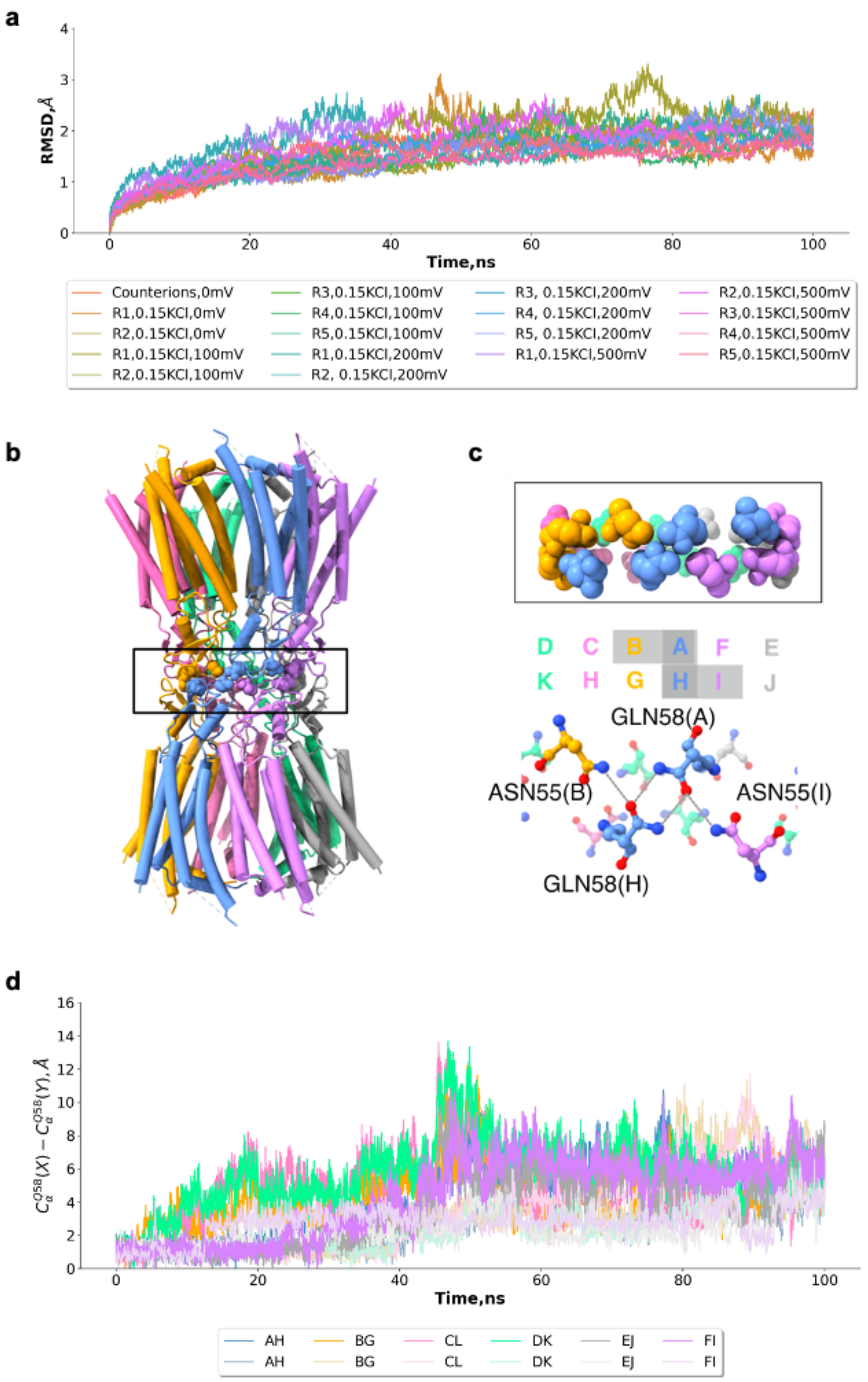


**Extended Data Figure 11. Molecular dynamics simulations of Cx43 GJC. a.** $C_\alpha$ root-mean-square-deviation from the cryo-EM structure during the different molecular dynamics simulations. **b.** Cx43 GJC structure depicted as cartoon with the interface region boxed. **c.** A zoomed-in view of the complex interface highlighted in A and the residues involved in the repeated motif are labelled highlighted as balls and sticks. **d.** Distance between the Q58 residues at the interface of the Cx43 during the two simulations at 0 mV and 0.15 KCl.

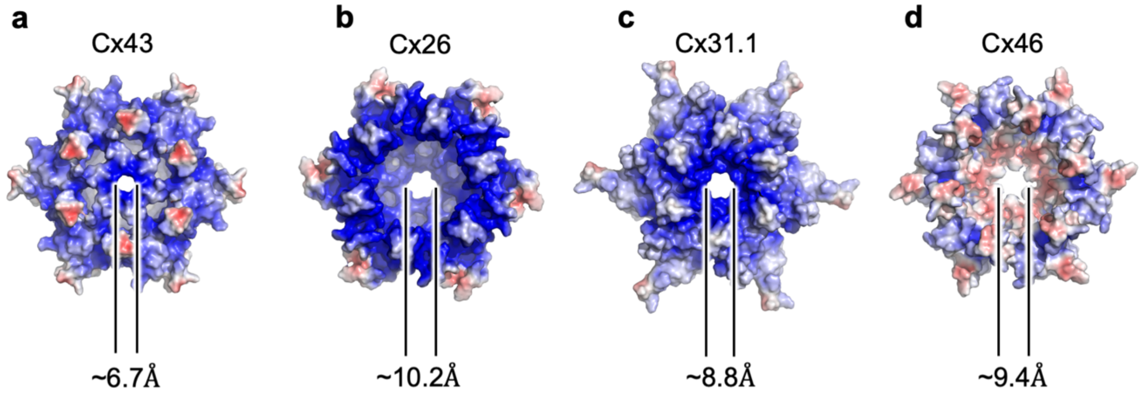


**Extended Data Figure 12. Electrostatic properties and dimensions of the gates in connexin channels. a-d.** The surface representations of the indicated connexin channels, Cx43, Cx26 GJC (PDB ID: 2zw3), Cx31.3 HC (PDB ID: 6l3t), Cx46 GJC (PDB ID: 7jkc), coloured according to the electrostatic potential, calculated using APBS [29]. The diameters of the gates were calculated using HOLE [30]. The images correspond to views of the channel from the cytosol, perpendicular to the membrane plane.

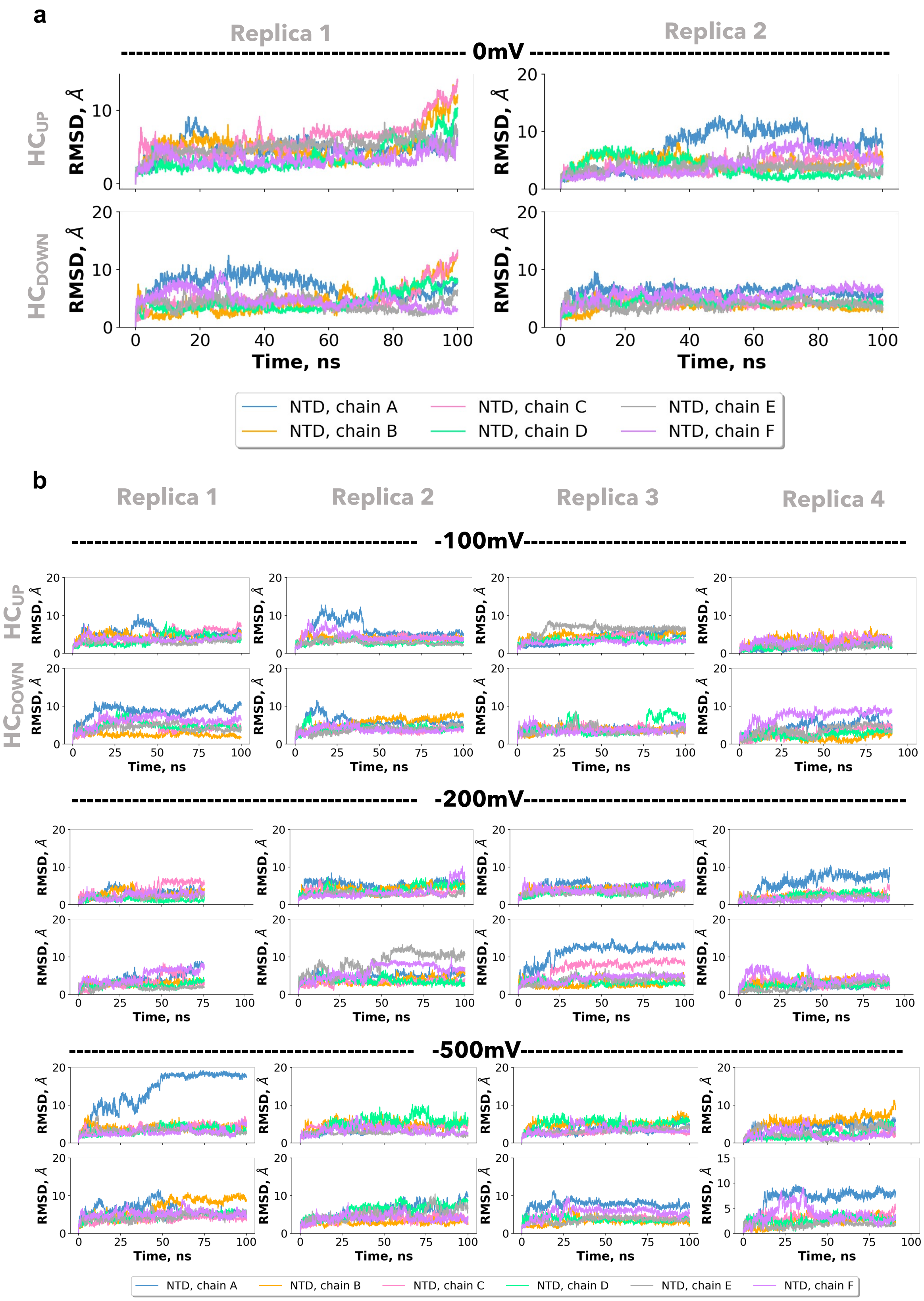


**Extended Data Figure 13. Molecular dynamics simulations of Cx43 GJC at different voltages.** Root-mean-square deviation (RMSD) of the NTD region for the 0mV replicas (**a**) and with increasing applied voltage replicas (**b**). The RMSD from the initial structure over time is depicted for the NTD region of every chain in the dodecameric structure. RMSD is shown for the different simulation replicates from left to right. Each panel includes the data for the chains forming the two hemichannels: A-F ($HC_{UP}$) and for chains G-L ($HC_{DOWN}$).

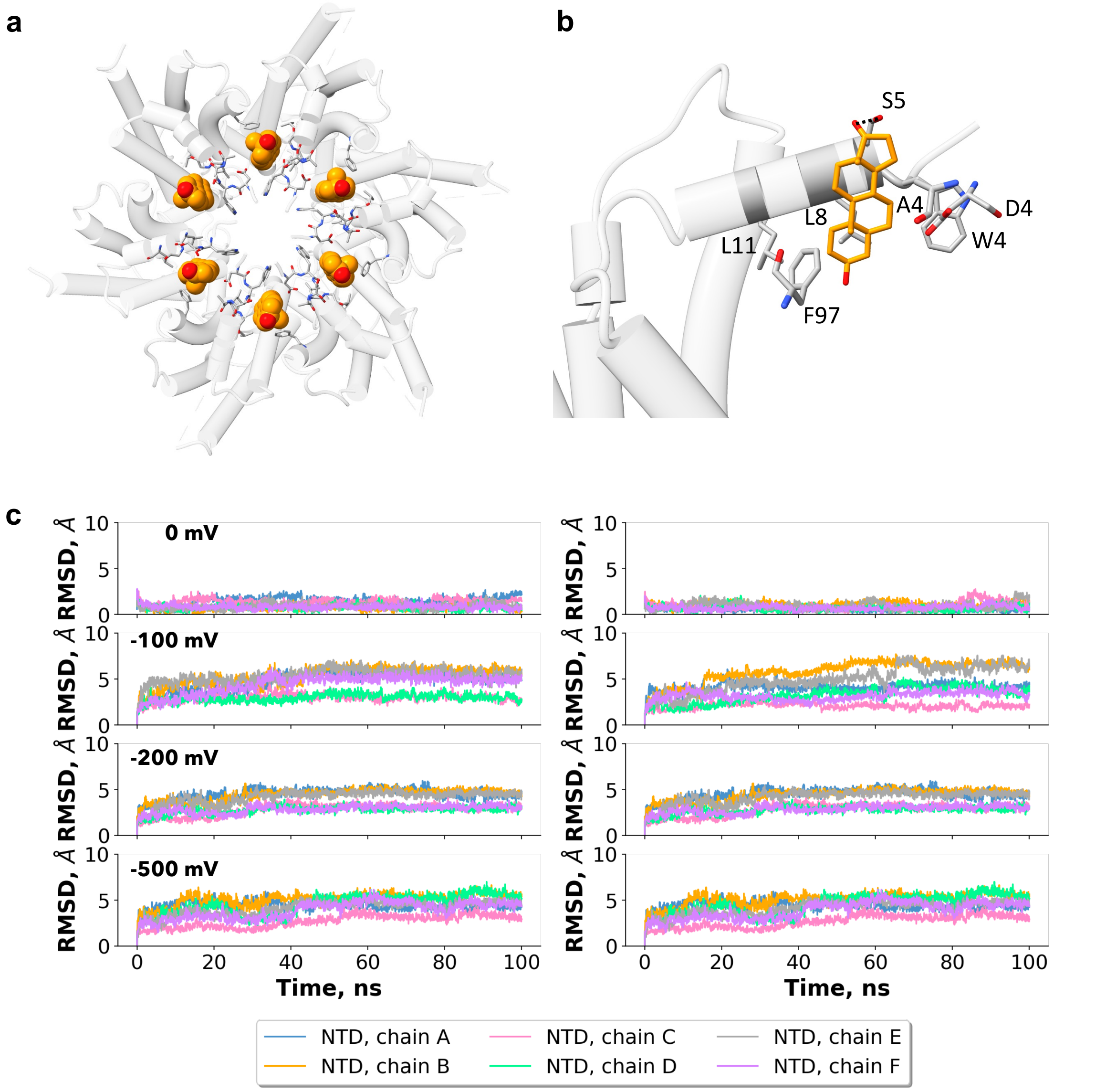


**Extended Data Figure 14. Molecular dynamics simulations of Cx43 HC with DHEA molecule in lipid-N site. a.** Cx43 HC structure depicted as a transparent cartoon with the DHEA molecule in orange as Van der Waals spheres and interacting residues from each chain as licorice. The MD simulations were performed with the HC model and a DHEA molecule as a lipid-N surrogate, with the DHEA and the NTD harmonically restrained (as described in Materials and Methods). **b.** A zoomed-in view of the NTD lipid site, with one DHEA molecule represented as licorice (orange) and the residues involved in contact with the molecule are highlighted as licorice and labelled accordingly. Only one chain is shown as a cartoon for clarity. **d.** Root-mean-square deviation (RMSD) of the NTD region for the different replicas (left to right) for the indicated applied voltages (top to bottom).

**Extended Data Table 1.** Cryo-EM data collection and image processing statistics

| Data collection | | | |
|---|---|---|---|
| Sample | Cx43 (Digitonin) | Cx43_nanodisc | |
| Instrument | FEI Titan Krios/Gatan K3 Summit/Quantum GIF | FEI Titan Krios/Gatan K3 Summit/Quantum GIF | |
| | | Cx43 GJC | Cx43 HC |
| Voltage | 300 | | |
| Electron dose($e^-$/Å) | 50 | | |
| Defocus range (μm) | -1 to -2 | | |
| Pixel size (Å) | 0.654 | | |
| Map resolution (Å) FSC threshold 0.143 | 2.26 | 2.95 | 3.98 |
| Map sharping b-factor (Å) | -49 | -86 | -100 |
| Number of particles | 50,471 | 10,886 | 20,526 |
| **Refinement** | | | |
| Model resolution (Å) FSC threshold 0.5 | 2.3 | 3.1 | 4.2 |
| Map CC | 0.84 | 0.84 | 0.77 |
| Model composition | | | |
| Protein residues | 2268 | 2268 | 1134 |
| ADP (B factor) | 31.90 | 40.02 | 158.95 |
| Bond length r.m.s.d. (Å) | 0.003 | 0.003 | 0.003 |
| Bond angle r.m.s.d. (°) | 0.552 | 0.514 | 0.678 |
| Validation | | | |
| MolProbity score | 1.47 | 1.37 | 1.79 |
| Clash score | 8.80 | 6.76 | 19.75 |
| Rotamer outliers (%) | 0 | 0 | 0 |
| Ramachandran plot | | | |
| Favored (%) | 98.92 | 98.24 | 98.92 |
| Allowed (%) | 1.08 | 1.76 | 1.08 |
| Disallowed (%) | 0 | 0 | 0 |

**Extended Data Table 2.** Ion permeation event statistics

| Applied transjunctional voltage | HC1 | | HC2 | | GJC | |
|---|---|---|---|---|---|---|
| | **K+** | **Cl-** | **K+** | **Cl-** | **K+** | **Cl-** |
| **0mV** | 176 ± 30 | 164 ± 56 | 397 ± 119 | 686 ± 122 | 0 | 0 |
| **100mV** | 275 ± 114 | 126 ± 33 | 490 ± 54 | 803 ± 232 | 1 | 0 |
| **200mV** | 382 ± 197 | 127 ± 32 | 409 ± 101 | 926 ± 136 | 0 | 0 |
| **500mV** | 243 ± 149 | 196 ± 115 | 628 ± 86 | 801 ± 270 | 1 | 1 |
